# Molecular insights into phycobilisome assembly pathway reveal crystalline bodies in cyanobacteria

**DOI:** 10.1101/2025.11.13.688327

**Authors:** Kelsey Dahlgren, Anton Avramov, Emily Koke, Akhil Gargey Iragavarapu, Andrew Hren, Francisco Javier Acosta-Reyes, Ayala Carl, Jonathan Herrmann, Liora Goldstein, John Spear, Jeffrey C. Cameron, Halil Aydin

**Affiliations:** Department of Biochemistry, University of Colorado, Boulder, CO, USA; Renewable and Sustainable Energy Institute, University of Colorado, Boulder, CO, USA; Interdisciplinary Quantitative Biology Program, BioFrontiers Institute, University of Colorado, Boulder, CO, USA; Department of Molecular Pathobiology, College of Dentistry, New York University, New York, NY, USA; Department of Chemical and Biological Engineering, University of Colorado, Boulder, CO, USA; Materials and Structural Analysis Division, Thermo Fisher Scientific, Hillsboro, OR, USA; Pain Research Center, New York University, New York, NY, USA; Department of Biochemistry and Molecular Pharmacology, Grossman School of Medicine, New York University, New York, NY, USA

**Keywords:** Cyanobacteria, photosynthesis, photosystem, phycobilisome, antenna, ApcE, light-harvest systems, thylakoid membrane, cryogenic electron tomography

## Abstract

In oxygenic photosynthetic organisms, light energy is converted to chemical energy to drive CO_2_ fixation reactions and sustain life on Earth. Cyanobacteria contain phycobilisome (PBS) complexes that play critical roles in light harvesting and directing light energy to the photosystem I and II reaction centers. The proper assembly of PBS components is an intricate process that is required for their activity and association with photosystem I and II. To understand the complex mechanisms regulating the PBS assembly, we knocked out the terminal emitter *apcE,* which forms the structural scaffold for the PBS core. *ApcE* knockout led to growth and pigment defects, including elevated levels of photosystem II and abnormal emission spectra. Light microscopy experiments revealed the accumulation of highly fluorescent puncta localized to the pole of *apcE* knockout cells. Further investigation using electron cryo-tomography identified highly repetitive crystalline arrays of densely packed PBS cylinders. Together, these data indicate that cyanobacteria may accumulate PBS components in the form of highly organized crystalline bodies as intermediates during PBS assembly.

## Introduction

Photosynthetic organisms evolved light-harvesting antenna to increase light absorption and funnel that energy towards photosynthesis. In plants and green algae, these antennae are called the light-harvesting complex, and are composed of chromophore-containing integral membrane proteins embedded in the thylakoid membrane around photosystem I (PSI) and photosystem II (PSII)^1^. In contrast, red algae and cyanobacteria contain large soluble and mobile antenna complexes called phycobilisomes (PBSs) that transfer energy to PSI and PSII^2–4^. PBSs are the source of red algae’s color and are particularly abundant in cyanobacteria, where they constitute up to 60% of soluble proteins in the cell^5^.

PBSs are composed of linker proteins and chromophore-containing phycobiliproteins (PBPs). In PBPs, light is absorbed by and transferred through covalently bound chromophores called phycobilins. The most common PBPs include phycoerythrin (PE), phycocyanin (PC), and allophycocyanin (APC). Phycoerythrobilin and/or phycourobilin are the phycobilin(s) in PE, which absorb light between 490-570 nm and fluoresce at 575 nm^5,6^. Phycocyanobilin is the phycobilin in PC and APC^6^, wherein the environment around the phycocyanobilin modifies its spectral properties in PC and APC. As a result, PC has a peak absorbance around 620 nm and emission at 640 nm, while APC has a peak absorbance around 650 nm and emission at 660 nm. ApcE and ApcD are specialized versions of APC that contain the terminal emitter phycocyanobilins with a further red-shifted emission between 675 and 680 nm and transfer energy to PSI and PSII^6,7^. At high enough concentrations, PBPs self-assemble into higher-order complexes of a 1:1 ratio of α:β subunits in solution^5^. The base unit (monomer) of PBPs is a heterodimer formed between the α and β subunits of a single PBP type. Three monomers come together to form a trimer, which takes a donut-like shape of a disc with a cavity in the center^2^. Two trimers can stack in a face-to-face manner to form a PC or PE hexamer^2,6^. However, linker proteins are needed to create larger structures from PBPs^2,6^.

A fully assembled PBS consists of rods radiating from a core with an arrangement of PBPs that allows light energy to flow downhill through the PBS towards the terminal emitters^8–13^. The PBS core is composed of several APC trimer stacks called cylinders. Tricylindrical PBS cores form a pyramid of two antiparallel cylinders on the bottom and a third cylinder on the top^9^. The cyanobacteria *Synechococcus sp.* PCC 7002 (PCC 7002) has a tricylindrical PBS core where each cylinder consists of four stacked APC trimer discs with aligned center cavities^9^. The bottom two cylinders of the pyramid have one edge of each APC trimer in the stack facing the thylakoid membrane^9^. ApcD and the chromophore-binding domain of ApcE replace one α APC subunit each, and ApcF replaces one β APC subunit on the thylakoid membrane side of each bottom cylinder^9^. Light energy flows from APC through the PBS core to the terminal emitters ApcE and ApcD, which directly transfer energy to PSII and PSI, respectively^2,14,15^. The ApcE protein is also essential for PBS assembly; it possesses several linker domains that fill the cavities in the center of APC trimers, creating APC cylinders of the proper length and lashing individual APC cylinders together into a fully assembled PBS core^5,9^. Additionally, removal of ApcE prevents the assembly of PBS cores, but not peripheral rods^16,17^. While tricylindrical PBS cores are common, the number of cylinders in a PBS core and the number of APC trimers in each cylinder are species-specific, depending on the number of linker domains in ApcE^2,18^. PBS cores also possess another linker protein called ApcC that occupies space on the end of PBS core cylinders and contributes to PBS stability^9,19^.

Fully assembled PBSs also contain peripheral rods made of rod-like stacks of PBP hexamers with linker proteins filling the central cavity of the PBP hexamers. The distal end of a PBS rod consists of the highest energy-absorbing chromophores, and the proximal end of a PBS rod contains PC to allow for downhill energy transfer from the PBS rod to APC, the PBS core^5^. The exact number of peripheral rods on a fully assembled PBS depends on the species and PBS core structure^2^. PBSs with tricylindrical cores typically have six PBS rods linked to the core with the rod-core linker protein CpcG^2^. The CpcG protein fills the cavity in the first PC hexamer of a peripheral rod has a motif that interacts with APC trimers in the PBS core^2^. Other rod linker proteins connect peripheral rod hexamers together^9^. A different version of PBSs called CpcL or rod-only PBSs lacks APC cores and is instead composed of a single peripheral rod with a CpcL linker protein taking the place of CpcG in the first hexamer^20^. CpcL modifies the environment around PC in the first hexamer to red-shift its emission and contains a transmembrane helix to associate the rod-only PBS with the thylakoid membrane^20,21^. While core-containing PBS preferentially transfer energy to PSII, rod-only PBS preferentially transfer energy to PSI. Therefore, the different PBS structures fulfill slightly different roles.

PBSs are involved in short-term and long-term adaptations to changing light conditions. In the short term, PBSs can provide photoprotection under high-light conditions. In cyanobacteria, an activated orange carotenoid protein associates with the PBS core to dissipate excess light energy as heat, which protects the photosynthetic reaction centers from photodamage^22^. PBSs are also involved in distributing energy between PSI and PSII on short time scales. The energy flux through PSI and PSII must be regulated to control the amount of ATP and reducing equivalents in the cell^23^. PBSs can associate or disassociate with PSI or PSII, depending on the needs of the cell, in a process called state transitions^23^. On longer time scales, the expression and structure of PBSs are modified in response to changing environmental conditions. For example, PBS expression and length of PBS peripheral rods are increased in low-light conditions to enhance the light-harvesting capacity^6,24^. Furthermore, some species of cyanobacteria and red algae undergo chromatic acclimation, in which the type of PBPs in peripheral rods is modified to optimize light energy harvesting under different wavelengths of light^24,25^. Cyanobacteria can also regulate their ratios of typical core-containing PBS to rod-only PBS. Expression of CpcL, the linker protein present in rod-only PBSs that preferentially associates with PSI, is increased in PSII-preferred green or orange light^20,26^. Therefore, CpcL may act to specifically increase the wavelength range of light energy transferred to PSI^3^.

Despite the physiological importance of PBS, there are unknowns within the assembly pathway of antenna megacomplexes. PBS formation begins with self-assembly of the PBPs into trimers and hexamers. Then, linker proteins stabilize the PBP trimers and hexamers into an arrangement that promotes energy transfer to the terminal emitters of the PBS. Yet, the sequence of events involving linker protein association with PBPs and bringing different parts of the PBS together into the fully assembled complex remains unclear. To better characterize the physiological changes that occur during PBS assembly, we knocked out *apcE* in the model cyanobacterium *Synechococcus* sp. PCC 7002. In addition to determining changes in growth rate, absorbance, and fluorescence characteristics, we observed stable, highly fluorescent puncta in the ApcE knockout with time-lapse fluorescence microscopy. The ApcE knockout was further characterized with cryo-electron tomography (cryo-ET) imaging, which revealed the highly fluorescent puncta to be crystalline protein assemblies composed of extremely long, tightly packed PBP cylinders. The crystalline accumulation of PBPs in the ApcE knockout provides insight into the assembly pathway of PBSs.

## Results

### Creation of ApcE knockout and knockdown strains

To knockout ApcE, we replaced the entire protein coding region of the *apcE* locus with a gentamycin resistance cassette using homologous recombination. ΔapcE has a growth defect and required many successive passages with antibiotic on 0.5% agar A+ plates to obtain a fully segregated strain (homologous for the gentamycin resistance cassette in the *apcE* locus). 0.5% agar was used instead of 1% because low agar percentages rescued growth of a different PBS knockout mutant^27^. Creation of the Δ*apcE* strain was verified by PCR of the *apcE* locus (Fig. 1A). To complement ApcE, we inserted a spectinomycin resistance cassette and the *apcE* protein coding region under control of a strong constitutive promoter (pK2) and into Neutral Site 1 (NS1) in the *ΔapcE*. Creation of the *ΔapcE+* strain was confirmed with PCR and sequencing of the apcE and NS1 loci (Fig. 1A).

**Figure 1.**
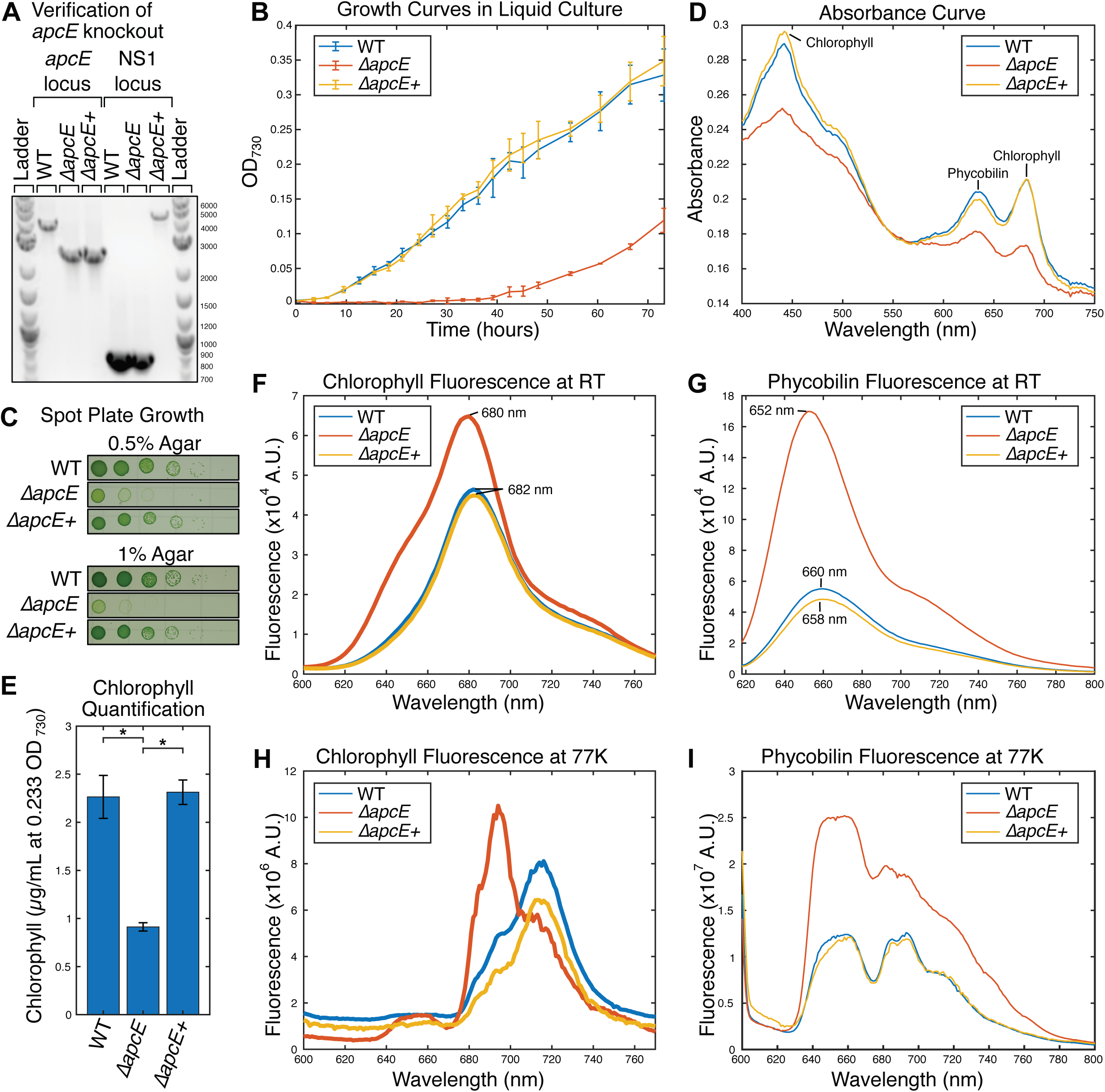
***ΔapcE* has growth and pigment defects** Experiments to characterize the growth and pigments of wild type PCC 7002 (WT), *ΔapcE*, and *ΔapcE+* were performed. (**A**) An electrophoresis gel of the PCR products from the *apcE* and NS1 loci. The sizes of the ladder bands are shown to the right of the gel in bp. The product of the WT *apcE* locus is 4.0 kb, while the *apcE* knockout has a size of 2.5 kb. The product of the WT NS1 site is 0.8 kb, while the *apcE* insertion in NS1 is 4.7 kb. The primers used are listed in table S1. (**B**) The growth was measured over time in liquid culture. OD_730_, a measure of cell density, was collected over 72 hours from cultures inoculated with 1 mL of culture diluted to the same OD_730_. The average OD_730_, a measure of cell number, of three replicates of each strain is displayed for each time point. The error bars represent one standard deviation. (**C**) The growth on 0.5 % (top) and 1 % (bottom) agar plates was compared. Cultures were diluted to the same OD_730_ and serial 1:10 dilutions were performed for each spot from left to right. A representative image from triplicate spot plates is displayed. (**D**) The absorbance spectra of each strain were compared. Liquid cultures were diluted to the same OD_730_ to normalize the cell number and absorbance spectra were collected. The panel displays the light wavelength on the x-axis and absorbance on the y-axis. Each line is an average of three replicates. (**E**) The amount of chlorophyll was quantified from each strain. In triplicate, cells from a liquid culture were diluted down to the same OD_730_, chlorophyll was methanol extracted, and the absorbance at 665 nm was measured and used to calculate the amount of chlorophyll per OD_730_. Error bars represent one standard deviation. (**F** and **G**) Chlorophyll (**F**) and phycobilin (**G**) emission spectra were collected at room temperature. Liquid cultures were diluted down to the same OD_730_ to normalize the cell number. The excitation wavelength was 440 nm for the chlorophyll emission spectra and 580 nm for the phycobilin emission spectra. The emission wavelength is on the x-axis and fluorescence is on the y-axis. Each line represents the mean of three samples. (**H** and **I**) Chlorophyll (**H**) and phycobilin (**I**) emission spectra were collected at 77 K. Each sample culture was concentrated to the same chlorophyll concentration in triplicate. The excitation wavelength was 440 nm for the chlorophyll emission spectra and 580 nm for the phycobilin emission spectra. The emission wavelength is on the x-axis and fluorescence is on the y-axis. Representative emission spectra from each strain are displayed.

### Δ*apcE* exhibits a growth defect

The growth rate of *ΔapcE*, WT, and *ΔapcE+* were monitored in liquid cultures and on plates with different agar percentages. While WT and *ΔapcE+* have similar growth, *ΔapcE* had a severe growth defect in both liquid cultures and on plates (Fig. 1, B and C). In liquid cultures *ΔapcE* has a longer lag phase and a slower growth rate than WT and *ΔapcE+*. On plates, *ΔapcE* displays a growth defect compared to WT and *ΔapcE+*, however this growth defect is more pronounced on 1% agar plates than 0.5% agar plates.

### *ΔapcE* displays a pigment defect

In both liquid cultures and on plates *ΔapcE* has a pigment defect by eye (Fig. 1C). To quantify pigment differences, liquid cultures of ΔapcE, WT, and ΔapcE+ were normalized to the same cell number (OD_730_ absorbance, OD_600_ cannot be used because 600 nm light is absorbed by pigments in cyanobacteria) and absorbance was measured across the visible spectrum (Fig. 1D). While WT and *ΔapcE+* have similar absorbance spectra, *ΔapcE* has decreased absorbance of chlorophyll (peaks at 440 and 680 nm) and phycobilin (peaks at 620 and 650 nm)^7^. Furthermore, the phycobilin to chlorophyll absorbance ratio (632/682nm) of 1.047 in Δ*apcE* is significantly higher (p<0.05) than the ratio 0.967 in WT or the ratio of 0.944 in Δ*apcE*^+^. To quantify chlorophyll, chlorophyll was methanol extracted from OD_730_ normalized number cells. Chlorophyll was significantly decreased in *ΔapcE* (p<0.05) to about 40% of WT or *ΔapcE+* (Fig. 1E).

### *ΔapcE* exhibits a high fluorescence phenotype with shifted peaks

To further characterize the pigments in *ΔapcE*, fluorescence spectra using both chlorophyll and phycobilin excitation wavelengths were collected at room temperature. Despite lower chlorophyll and phycobilin absorbances than WT and ΔapcE+, *ΔapcE* has higher fluorescence emissions when excited by chlorophyll and especially phycobilin wavelengths (Fig. 1, F and G). Furthermore, the chlorophyll fluorescence peak is shifted to 680 nm in Δ*apcE* from 682 nm in WT and Δ*apcE*^+^ and the phycobilin fluorescence peak is shifted to 652 nm in Δ*apcE* from 660 nm in WT and 658 nm in Δ*apcE*^+^.

To characterize the fluorescence phenotype in greater detail, fluorescence spectra were also collected at 77K. At this temperature, the biochemical and physiological processes that modulate fluorescence are inhibited, allowing for an accurate measurement of the fluorescent molecules^28^. Additionally, the fluorescence emission spectra of the PBPs, PSI, and PSII can be easily distinguished at 77K^28^. Furthermore, at this temperature light energy can be absorbed and transferred to lower energy state pigments, allowing the connection between PBSs and PSI or PSII to be interrogated^29^. Excitation of chlorophyll at 440 nm results in two major emission peaks, one from PSII split between 685 and 695 nm, and one from PSI at 720 nm^28,30^. *ΔapcE* has an increased ratio of PSII/PSI compared to WT and *ΔapcE+* (Fig. 1H). *ΔapcE* 77K 440nm emission spectra also has a small hump around 650-660 nm likely originating from the increased PBP fluorescence in Δ*apcE*.

At 77K, excitation of phycobilin at 580 nm results in several emission peaks. As in the chlorophyll emission spectra, there are peaks from PSII at 685/695 and PSI at 720 nm (Fig. 1I). These peaks are the product of PSII- and PSI- coupled PBSs that absorb light energy and transfer it to their respective photosynthetic reaction centers^28^. There is also a “free PBS” peak at 640-660 nm originating from uncoupled PBSs that do not transfer light energy they absorb to PSI or PSII and partially assembled or unassembled PC (640 and 650 nm) and APC (660 nm)^7,23^. At 77K, the increased ratio of free PBS to PBSs coupled to PSII or PSI in *ΔapcE* compared to WT or *ΔapcE+* is clearly observed, demonstrating impaired energy transfer from PBSs in *ΔapcE*.

### ApcE CRISPRi knockdown has similar, but less severe phenotypes than ΔapcE

CRISPRi was used to create an inducible ApcE knockdown strain (fig. S1). In PCC 7002 containing dCas9 under control of anhydrotetracycline (aTC) in the *ascA* locus, a gRNA spacer targeting the promoter of *apcE* under control of IPTG was inserted into the *glpK* locus using a kanamycin cassette for selection. The apcE CRISPRi strain was confirmed with PCR and sequencing of the *glpK* locus. The NT (non-targeting) CRISPRi containing a gRNA spacer lacking significant homology to any site in the PCC 7002 genome was also created and confirmed to act as a control for the apcE CRISPRi strain. For CRISPRi to be active, both aTC and IPTG are added to the CRISPRi strain culture to induce expression of dCas9 and the gRNA, which associate and bind the DNA targeted by the gRNA, blocking transcription of the targeted gene (fig. S1). ApcE CRISPRi induced with aTC and IPTG had defects in growth, pigments, and fluorescence compared to uninduced ApcE CRISPRi or NT CRISPRi induced with aTC and IPTG (fig. S1). The defects observed in the ApcE CRISPRi knockdown were not as severe as those observed in *ΔapcE*.

### *ΔapcE* has fluorescent puncta

*ΔapcE* was examined on a time-lapse fluorescence microscope to visualize the spatial patterns of different autofluorescent pigments in the cells (Fig. 2A). In WT cells, endogenous fluorescence tends to be relativly evenly distributed across the cells with slightly higher fluorescence just inside the edges of the cell corresponding to the thylakoid membrane. In contrast, extremely bright puncta were observed in the Cy5 channel of Δ*apcE* images (ex: 640 nm, em: 665-715 nm). The Cy5 excitation wavelength is near the center of the phycobilin absorption peak and on the very high energy range of the red chlorophyll absorption peak (fig. S2). The emission range of the Cy5 filter includes the low energy range of phycobilin fluorescence and the majority of chlorophyll fluorescence. These puncta in Δ*apcE* are also observed in the RFP channel (ex: 555 nm, em: 571-628 nm). The RFP channel is phycobilin specific; Both the excitation and emission wavelengths of the channel exclude chlorophyll excitation and emission through their placement on the extremely high energy edge of phycobilin fluorescence excitation and emission ranges (fig. S2). While Δ*apcE* has highly fluorescent puncta, normalized lookup tables allowed visualization of the autofluorescence signal distributed across the WT and Δ*apcE*^+^ cells to also be present in Δ*apcE* cells. (Fig. 2A). To quantify the puncta, the fluorescence values of a line drawn between the cell poles were collected (Fig. 2B) and the “puncta value” was calculated by dividing the maximum fluorescence by the first quartile fluorescence (Fig. 2, E and F). The puncta value was used instead of raw maximum fluorescence values to normalize for any uneven illumination across the field of view on the microscope. Cells were defined as possessing fluorescent puncta if their puncta value was greater than three standard deviations above the WT average. One percent of WT cells have fluorescent puncta in both the Cy5 and RFP channels. In the Cy5 channel, 94% of Δ*apcE* cells have puncta. In the RFP channel, 88% of ΔapcE cells have puncta. Δ*apcE*^+^ has puncta in 4% of cells in the Cy5 channel and 5% of cells in the RFP channel. The presence of puncta is mainly driven by higher fluorescence maximum values in Δ*apcE* (fig. S2). Interestingly, the average first quartile fluorescence value is decreased in Δ*apcE* compared to WT and Δ*apcE*^+^ in the Cy5 channel, but the reverse is true in the RFP channel (fig S2). This discrepancy may explain why a lower proportion of Δ*apcE* cells have puncta in the RFP channel than the Cy5 channel.

**Figure 2.**
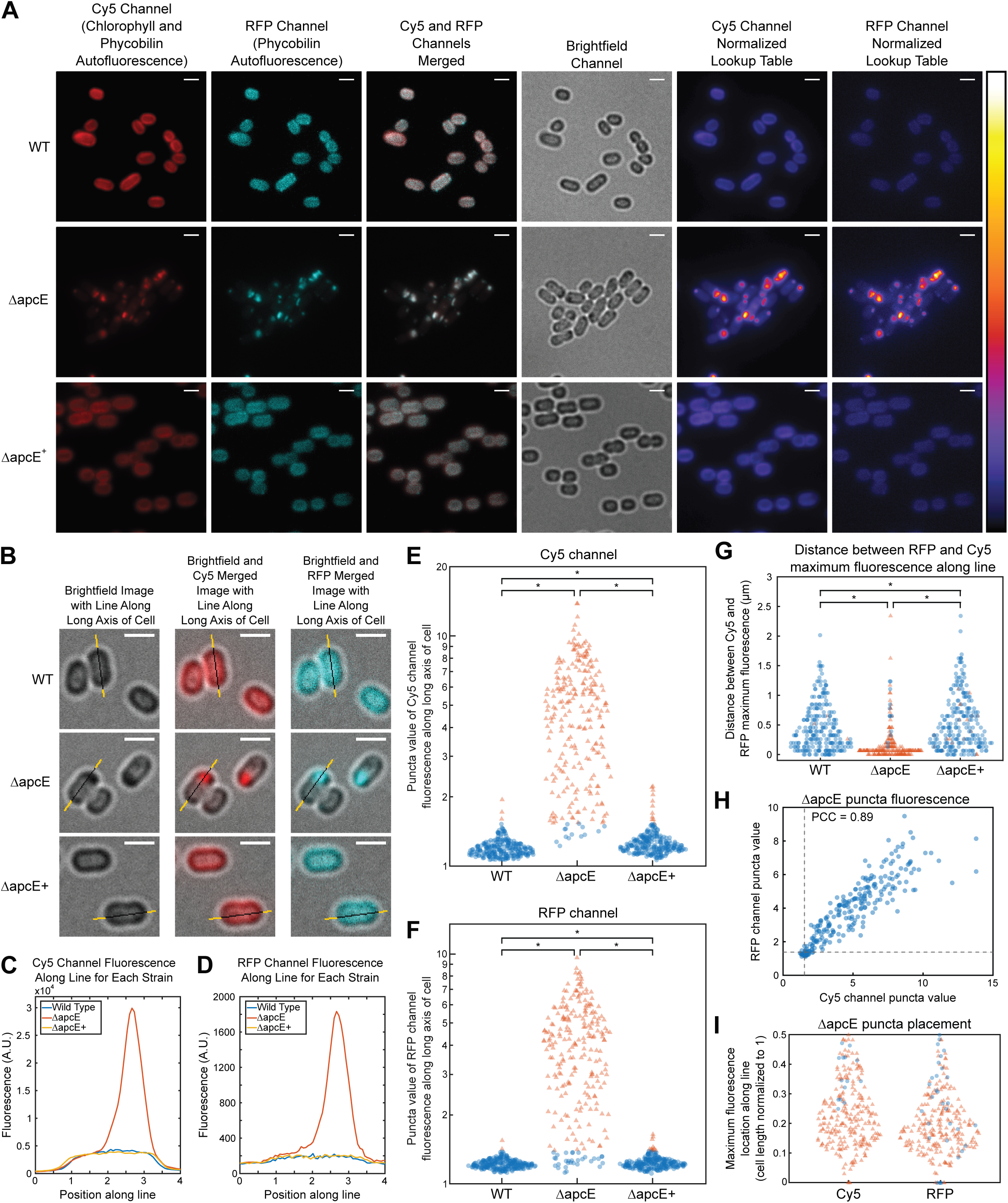
Δ*apcE* has fluorescent puncta. Fluorescence microscopy was performed to analyze the subcellular localization of autofluorescent pigments in PCC 7002. (**A**) Fluorescence microscopy images of WT (top row), Δ*apcE* (middle row), and Δ*apcE+* (bottom row). Scale bars representing 2 µm are present in the top right corner of each image. The first column displays Cy5 channel fluorescence (ex: 640 nm em: 665-715). The second column displays RFP channel fluorescence (ex: 555 nm em: 571-628). The third column displays a merged image of both the Cy5 (red) and RFP (cyan) channels. The fourth column displays transmitted light. The fifth and sixth columns display the Cy5 and RFP channels with normalized lookup tables between WT, Δ*apcE*, and Δ*apcE+*. The scale for the lookup tables for the fifth and sixth columns is on the far right. (**B**) Close-up images of WT (top row), Δ*apcE* (middle row), and Δ*apcE+* (bottom row). The top right corner of each image contains a scale bar representing 2 µm. The first column displays transmitted light images, the second column displays a merged transmitted light and Cy5 channel image, and the third column displays a merged transmitted light and RFP channel image. In each image, a 4 µm line is drawn through a single cell. (**C**) Cy5 and (**D**) RFP fluorescence values of the lines drawn through cells WT (blue), Δ*apcE* (red), and Δ*apcE+* (yellow) in Fig. 2B are plotted against their position in the line. The distribution of puncta values (the ratio of maximum fluorescence value to the first quartile fluorescence value along a pole-to-pole line) in the (**E**) Cy5 and (**F**) RFP channels were plotted on log scales for WT (left), Δ*apcE* (middle), and Δ*apcE+* (right). Red triangles represent values greater than 3 standard deviations more than the WT puncta value average. The remaining values are displayed as blue circles. Significantly differences (p<0.001) are marked with asterisk (*). 250 replicates are plotted for each strain. (**G**) The distance between the position of the maximum fluorescence value of the RFP and Cy5 channels along a line drawn between cell poles for WT (left), Δ*apcE* (middle), and Δ*apcE+* (right). Cells with puncta values greater than 3 standard deviations above WT puncta value in both the Cy5 and RFP channels are plotted as red triangles and all other cells are plotted as blue circles. 250 replicates are plotted for each strain. (**H**) The correlation between puncta value from the RFP and Cy5 channels for Δ*apcE* cells is displayed. The dotted lines represent 3 standard deviations above the WT puncta value. The Pearson correlation coefficient (PCC) value is displayed. (**I**) The position of the maximum fluorescence value within Δ*apcE* cells is displayed. Each cell was normalized to a length of 1, and the distance between the maximum fluorescence value in the Cy5 (left) and RFP (right) and the nearest cell pole was calculated for 250 replicates. Cells with puncta values greater than 3 standard deviations above WT puncta value are plotted as red triangles and all other cells are plotted as blue circles.

There is a correlation between puncta location in the Cy5 and RFP channels in Δ*apcE* (Fig. 2A). The distance between the positions of Cy5 and RFP maximum fluorescence on pole-to-pole line is significantly (p<0.05) lower in Δ*apcE* than WT or Δ*apcE*^+^ (Fig. 2G). Furthermore, there is a higher correlation (Pearson Correlation Coefficient (PCC) = 0.89) (Fig. 2H) between the puncta values of the Cy5 and RFP channels in Δ*apcE* compared to WT (PCC = 0.39) or Δ*apcE*^+^ (PCC = 0.50). Therefore, fluorescent puncta observed in the Cy5 and RFP channels of Δ*apcE* images are likely imaging the same phenomena. Due to the fluorescent properties of the puncta in both the Cy5 and RFP channels, the puncta likely originate from a phycobilin-containing cellular structure or aggregate. The puncta tend to be located away from the center of the cell and closer to cell poles (Fig. 2I).

### Time-lapse imaging reveals dynamics of highly fluorescent puncta

Time-lapse imaging was used to observe the dynamics of the fluorescent puncta in Δ*apcE* (Fig. 3). The Δ*apcE* fluorescent puncta are stable structures. The puncta present at or near the start of time-lapse imaging can be tracked for over 90 hours across up to 6 cell divisions (movies S1 and S3, filled arrows). Only cells that ceased growing and dividing showed complete dissipation of existing fluorescent puncta (movies S2 and S3, open arrows). In these cases, the length of time for puncta fluorescence loss after cell growth stopped was variable, from less than 20 hours (movie S2) to greater than 50 hours (movie S3). Growth was observed in both cells with and without fluorescent puncta, so fluorescent puncta are not required for cell growth in Δ*apcE*. However, growing Δ*apcE* cells developed puncta within a few hours after division if they did not inherit one (Fig. 3A). The time to develop puncta is less than the time between cell divisions, so cells possess fluorescent puncta by the time they divide (movies S1 and S3). The majority of puncta are inherited by only one of the daughter cells, although we also observed puncta that were split between daughter cells (Fig. 3B).

**Figure 3.**
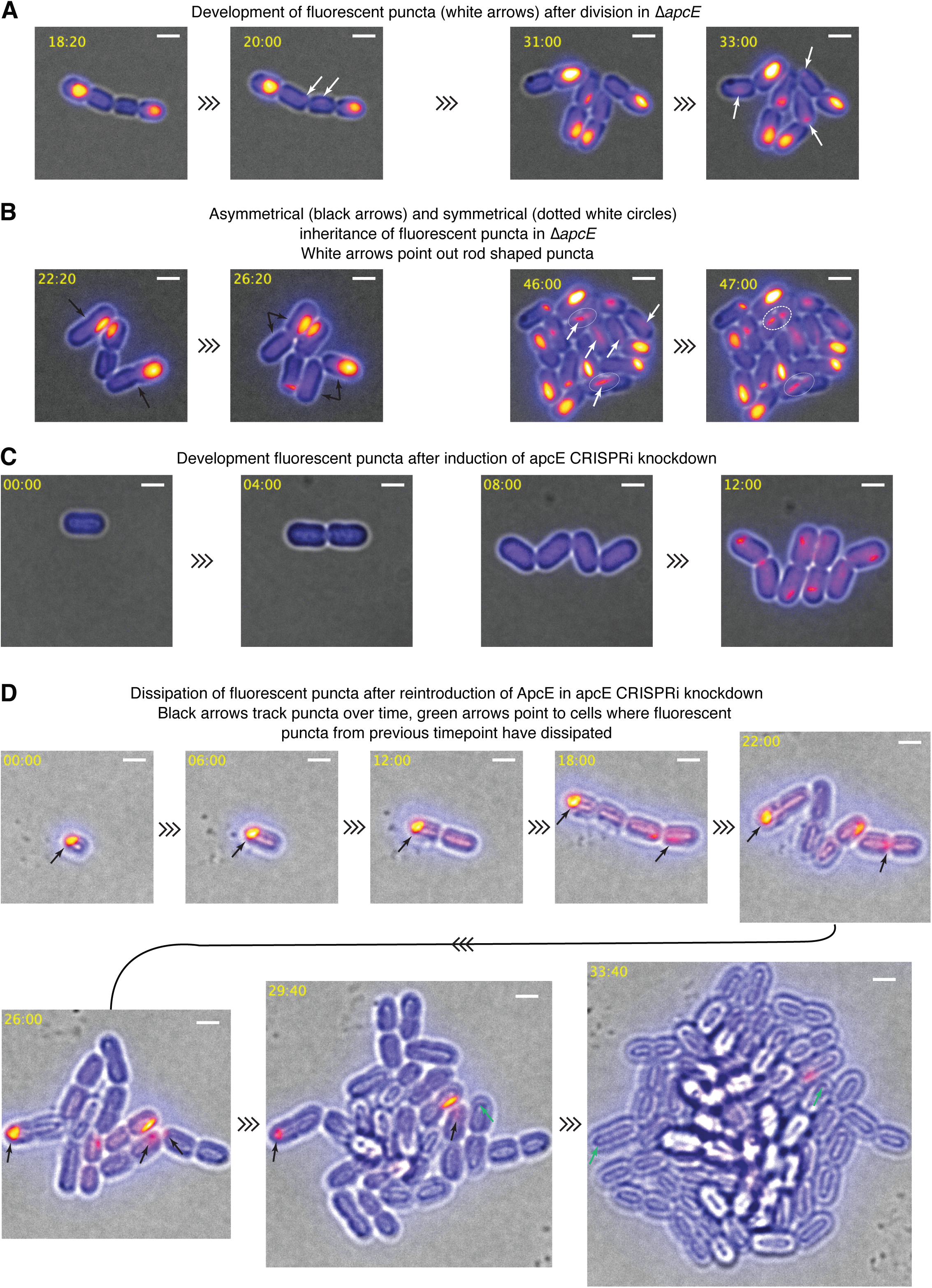
Time-lapse microscopy of Δ*apcE* and *apcE* CRISPRi knockdown. Time-lapse fluorescence microscopy was performed on Δ*apcE* and the *apcE* CRISPRi knockdown to characterize the dynamics of the highly fluorescent puncta. In all image panels, the time point is displayed on the top left and a 2 µm scale bar is displayed in the top right. All image panels are merged images of transmitted light (black and white) and the Cy5 channel (scale in figure 2A). (**A**) Still images from movie S1 are displayed to show puncta development after division in Δ*apcE*. Each image set displays several newly divided cells without fluorescent puncta in the left image and white arrows pointing to newly developed puncta in the right image taken several hours later. (**B**) Stills from time-lapse imaging of Δ*apcE* showing symmetrical and asymmetrical inheritance of fluorescent puncta. Each set of two images displays the same colony at two time points. The image set on the left are from movie S3 and the image sets on the right are from movie S1. (**C**) Stills from movie S4 of a time-lapse of induction of the *apcE* CRISPRi knockdown. The *apcE* CRISPRi strain was precultured in the absence of IPTG and aTC so *apcE* expression was not inhibited. Knockdown of *apcE* was induced by the presence of IPTG and aTC in the agarose pad used as a growth substrate during time-lapse imaging. (**D**) Stills from movie S6 of a time-lapse of recovery of *apcE* expression in the *apcE* CRISPRi knockdown strain. Cells were precultured with aTC and IPTG to ensure *apcE* knockdown before time-lapse imaging. The agarose pad used as a growth substrate during time-lapse imaging did not contain aTC or IPTG, which allowed *apcE* expression to recover. Black arrows point to puncta that were tracked over time. Green arrows point to cells where the tracked highly fluorescent puncta (black arrows) have faded away.

The *apcE* CRISPRi knockdown was also utilized to observe the dynamics of the highly fluorescent puncta. First, puncta formation was observed in when CRISPRi was induced in the *apcE* CRISPRi knockdown strain to prevent *apcE* expression. Cells were precultured in the absence of inducers to ensure that ApcE was present when time-lapse imaging began. However, inducers were present in the agarose pad covering cells during time-lapse imaging, so *apcE* repression began when time-lapse imaging began. Highly fluorescent puncta formed after about 10 hours in these conditions (Fig. 3C and movies S4 and S5). The puncta persisted over multiple cell divisions, similar to our observations of Δ*apcE*. We also observed reintroduction of ApcE into an induced *apcE* CRISPRi knockdown. Cells were precultured with aTC and IPTG to prevent *apcE* expression. Then cells were washed and placed onto agarose pads lacking aTC and IPTG to allow *apcE* expression to resume during time-lapse imaging. We observed fluorescent puncta in cells at the start of time-lapse imaging. In the first ∼20 hours, cells that did not inherit fluorescent puncta produced fluorescent puncta similar to Δ*apcE*. However, after ∼25 hours, cells that did not inherit a fluorescent punctum ceased production of fluorescent puncta while continuing to grow and divide (Fig. 3D and movies S6 and S7). Furthermore, fluorescent puncta that were present at the start or formed at the beginning of time lapse imaging faded away over time. In contrast to the fluorescent puncta dissipation observed in Δ*apcE*, the fluorescent puncta dissipation during ApcE reintroduction occurred in growing cells (movies S6 and S7).

### Visualization of *ΔapcE* fluorescent puncta by cryo-collerative light and electron microscopy

To determine the molecular origins of the *ΔapcE* fluorescent puncta, we turned to on-grid cryo-correlative light and electron microscopy (cryo-CLEM) experiments. Freshly grown cyanobacteria were applied onto EM grids and plunge frozen into liquid ethane using Leica EM GP2. However, cells vitrified on EM grids cannot be directly imaged by electron cryo-tomography (cryo-ET) due to the thickness of the specimen. To facilitate cryo-ET imaging, we utilized cryo-focused ion beam (cryo-FIB) milling to thin cyanobacterial samples into cellular sections (lamellae), which are ∼150-250 nm in thickness and transparent to the electron beam. We first employed cryo-correlative light microscopy imaging to identify regions positive for *ΔapcE* fluorescent puncta on frozen-hydrated grids by illuminating cyanobacterial cells at an excitation wavelength of 560 nm, which excites phycobilin. Consistent with the fluorescence microscopy imaging, Δ*apcE* cells frozen on EM grids exhibited abundant bright fluorescent puncta (Fig. 4A). These light microscopy images were then correlated with scanning electron microscopy (SEM) data to determine target areas for lamella preparation (Fig. 4, A to D). After cryo-FIB milling, accurate correlation between phycobilin fluorescent signal and vitrified PCC 7002 was verified with low magnification SEM imaging of the lamellae surfaces, revealing intact cyanobacterial cells containing thylakoid membrane networks positioned near the plasma membrane (Fig. 4D). Together, these experiments demonstrated that cryo-ET imaging could reliably provide a detailed view of the Δ*apcE* fluorescent puncta in near-native conditions.

**Figure 4.**
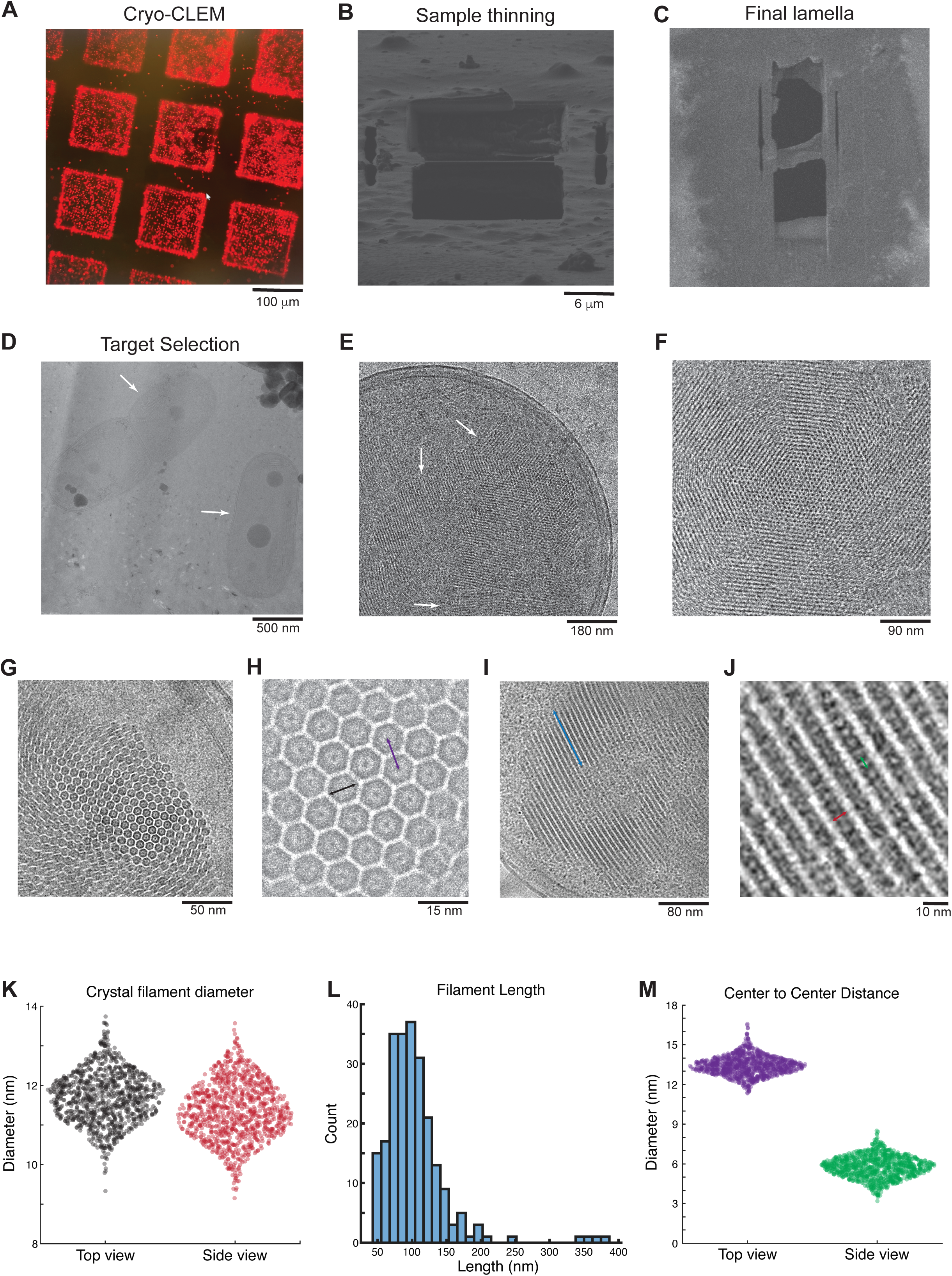
In situ cryo-ET workflow from sample preparation to image analysis. (**A**) Cryo-CLEM image shows the distribution of the chlorophyll fluorescence emitted from the Δ*apcE* cells. Scale bar, 100 µm. (**B**) A representative cryo-SEM image of the samples thinned to approximately 150-200 nm. (**C**) A representative image of the cryo-FIB milled sample showing the final lamella after polishing. (**D**) Montage overview of the lamella displays intact frozen, hydrated cyanobacterial cells. White arrows indicate the position of the cells. (**E** and **F**) A zoomed in image of the Δ*apcE* cell crystalline arrays. Scale bars are 500 nm (**D**), 180 nm (**E**), 90 nm (**F**). (**G**) A tomogram reconstructed from tilt series collected on lamella reveals the presence of crystalline arrays occupying the cytosolic space of the Δ*apcE* cell. (**H**) Representative image of the 2D crystalline array in the top view, showing hexameric structures Scale bars are 50 nm (**G**) and 15 nm (**H**). (**I** and **J**) Side view images of the crystal lattice display the assembly of rod-shaped molecules with varying lengths. Scale bars are 80 nm (**I**) and 10 nm (**J**). (**K**) Quantification of crystalline lattice subunit diameter (black arrow in **H** and red arrow in **J**) was performed by measuring the surface area of each subunit from top and side view images. A total of ∼1000 subunits were analyzed from different tomogram reconstructions. (**L**) Rod lengths were quantified from side view images and are shown as a bar graph (blue arrow in **I**). A total of ∼1000 rods were analyzed from different tomogram reconstructions. (**M**) Quantification of inter and intra particle distance from top and side view images (purple arrow in **H** and green arrow in **J**). A total of ∼1000 hexamers were analyzed from different tomogram reconstructions.

### Cryo-ET of FIB-milled cyanobacteria reveals a 2D crystalline lattice

Cryo-FIB milled lamellae were imaged using a Krios cryo-transmission electron microscope (cryo-TEM) operated at 300 KeV and equipped with a Selectris energy filter and Falcon 4i direct electron detector. The tilt series of the targets was acquired with Tomography 5 Software using a dose-symmetric tilt-scheme^31^, starting at a pre-tilt angle defined by the milling angle and ranging ±55° around it in 3° increments. In total, we obtained 22 tomograms of cryo-FIB-milled samples, which showed dramatic improvements in image quality and contrast compared to our previous unthinned samples, allowing us to investigate the identity of *ΔapcE* fluorescent puncta. The tomograms were reconstructed from tilt series with IMOD^32^. Visual inspection of the aligned tomograms revealed the characteristic ultrastructure of cyanobacterial plasma membranes and thylakoid networks inside cells. Further analyses of the Δ*apcE* tomograms also revealed large proteinaceous lattices with a mean area of 0.242 ± 0.080 µm² in the cytoplasmic space (Fig. 4, E and F). These crystalline structures observed by cryo-ET consist of well-defined subunits that are highly repetitive with an apparent p6 symmetry. The lattice subunits show similar structural features and conformational states, leading to the formation of compositionally homogeneous crystalline arrays within *ΔapcE* cells (Fig. 4, G to J). While the tomograms capturing the top view of the lattice clearly demonstrated the dense packing of hexameric structures (Fig. 4H), the side view tomograms revealed that each subunit adopts a rod shape, suggesting that the alignment and tight packing of these elongated molecules in the same directions creates the crystalline array (Fig. 4J). When we analyzed the subunit diameters by taking three diameter measurements of each subunit in the XY plane and averaging them, we found that the mean diameter of each subunit is 11.70 ± 0.64 nm (Fig. 4K). In the Z plane, the distribution of subunit length varied significantly, ranging from ∼40 to 400 nm with a mean rod length of 104.18 ± 47.47 nm (Fig. 4L). Finally, the center-to-center mean inter-particle distance was measured around 13.14 ± 0.90 nm within the crystalline lattice (Fig. 4M). These data strongly suggest that the assembly of rod-shaped molecules into the crystalline array through their long axis allows the dense packing of subunits while facilitating the incorporation of rods with varying lengths.

### Crystalline arrays are formed by PBS rods in *ΔapcE* cells

The large size and repetitive appearance of the crystalline arrays in *ΔapcE* cells suggested that relatively abundant cyanobacterial proteins may form these assemblies. Cryo-ET tomograms of the crystalline arrays revealed structural features that resemble the light-harvesting (PBS) rods, which can account for up to 60% of the total soluble protein mass in cyanobacteria^5^. To determine the potential molecular identities of crystalline lattice subunits, we sought to examine the molecular morphologies of PBS using three-dimensional (3D) models. We performed detailed structural analysis comparing the structures observed in Δ*apcE* tomograms and PBS structures from PCC 7002 and other organisms. PBS peripheral rods are formed by the stacking of several disc-shaped hexamers on one another in a back-to-back fashion and can span tens of nm longitudinally. The PBS hexamers have an average diameter of ∼11 nm and a height of ∼6 nm in various organisms^33^. Visual inspection of the cryo-ET tomograms exhibited the stacking of disk-shaped molecules through the long axis of the crystalline lattice subunits (Fig. 4, G to M). These assemblies typically contained more than 20 disk-shaped molecules and reached an average length of 100-150 nm (Fig. 4, I to M). Our measurements revealed that the dimensions of the densities for individual disk-shaped molecules and the assembled rods in tomograms match the tertiary structures of hexamers and PBS rods, indicating that the elongated densities of the lattice could be PBS rods (Fig. 4). Additionally, we did not observe any interactions between the crystalline array and thylakoid or plasma membranes in Δ*apcE* tomograms, which is consistent with the lack of the PBS core-membrane linker ApcE. Together, these analyses suggest that in the absence of ApcE, PBS peripheral rods fail to attach to a PBS core and instead nucleate to form a highly fluorescent dense array of rods in the cytosolic space of cyanobacteria.

### In-cell structure of PBS rods

To further elucidate the content of crystalline lattice subunits, we performed additional structural analysis in 3D. We first generated a lattice correlation map of the crystalline array using the Δ*apcE* tomograms (Fig. 5A). The 3D map provided a glimpse into the architecture and the assembly mechanism of the lattice. While the top view of the crystalline array showed hexameric molecules forming a predominantly p6 symmetric lattice, the side view of the map revealed variations in rod length (Fig. 5A). In cyanobacteria and red algae, changes in local light and environmental conditions and dynamic assembly mechanism of these complexes result in the formation of highly variable rod structures^2^. Given that the packing of the rods is mediated through lateral interactions in the lattice, the *ΔapcE* cells could accommodate the incorporation of rods with different lengths into a symmetric lattice and promote the nucleation and lateral growth of the crystalline assemblies *in vivo*. To further support the hypothesis that the crystalline lattice is formed by PBS rods, we conducted subtomogram analysis on individual subunits of the assembly. We semi-automatically picked 32907 lattice subunits from our entire dataset. These subtomograms were processed without symmetry and showed a near homogeneous population of well-defined cylindrical assemblies. Through 3D refinement and classification in RELION^34^, the final class of subtomograms revealed a rod-shaped structure resolved to approximately 19 Angstroms (Fig. 5B and fig S3). At this resolution, the cryo-ET map displayed cylindrical stacks of disk-shaped substructures that resemble PBP hexamers^9^ (Fig. 5, B and C). We observed weak densities connecting the lattice subunits between columns, suggesting that non-covalent interactions primarily mediate the interconnectivity of the crystalline array. PBS peripheral rod hexamers from PCC 7002 were fitted into the density map of crystalline lattice using a recently reported cryo-EM structure^9^ as reference model, showing consistency and good agreement with the observed densities (Fig. 5B). We further performed real space refinement using the *in situ* cryo-ET map and PBS models and demonstrated that the refined models fit well into the cryo-ET map with a map-model correlation coefficient of 0.58 (main chain) (Fig. 5, B and C). These analyses, along with the agreement between our measurements of assembled lattice subunits in 2D, indicate that the subunit arrangement in our models is an accurate representation of the PBS rod lattices formed in Δ*apcE* cells *in vivo*.

**Figure 5.**
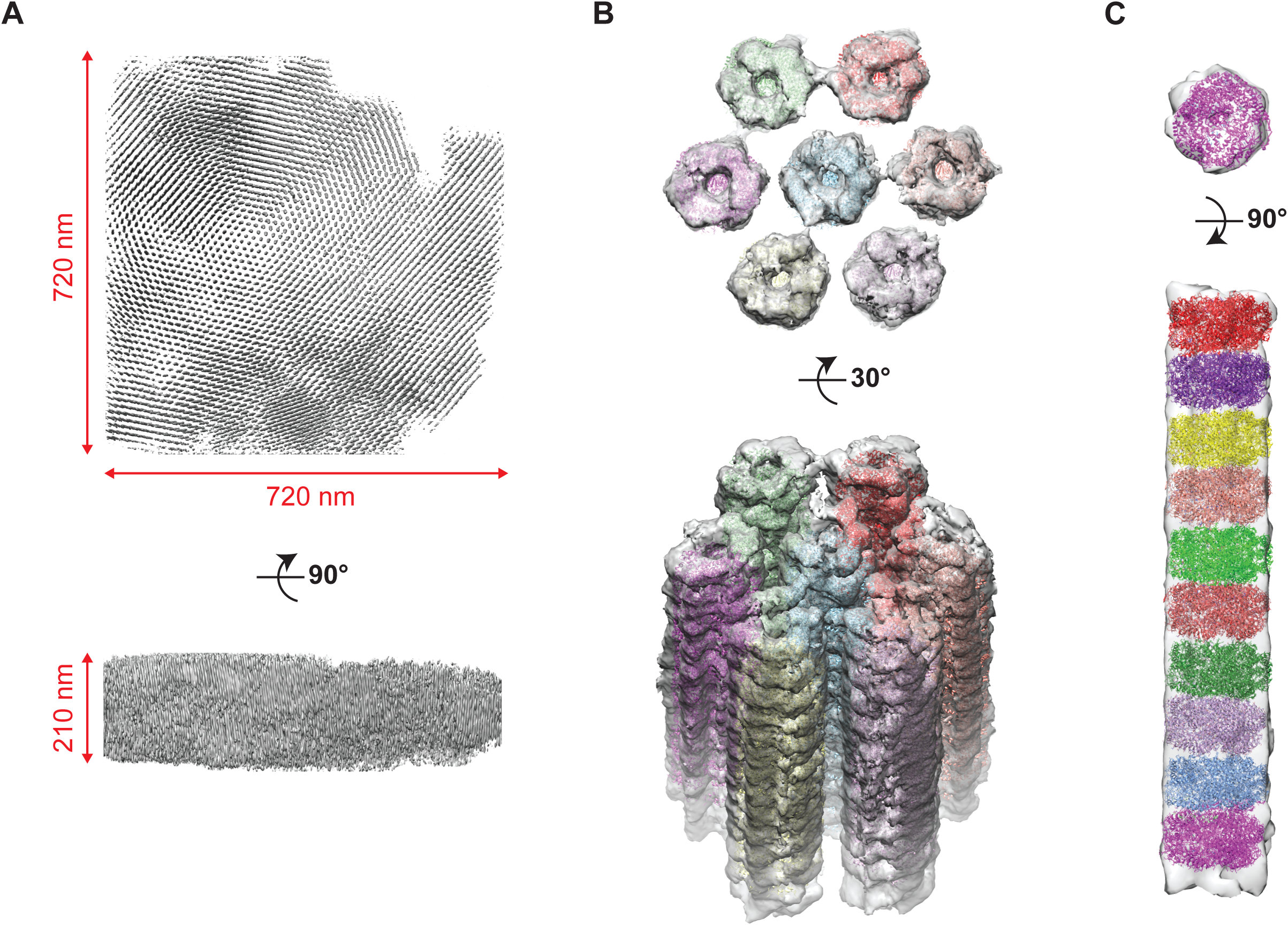
Figure 5. Overall structure of the PBS crystalline array. (**A**) 3D lattice correlation map of the crystalline array illustrates organized and dense packing of PBS rods with various lengths. Top view of the lattice reveals minimal contacts between lattice subunits and highlights the p6 symmetry. (**B**) Subtomogram averaging reconstruction of PBS arrays, showing in the top and tilted views. Peripheral rod models of the PBS from *Synechococcus sp.* PCC 7002 were extracted from the structure and fitted into the cryo-ET density map (surface representation). Atomic models of PBS peripheral rods are colored differently for each subunit of the crystalline array. (**C**) Top and side views of a single subunit density show strong agreement with the PBS hexamer models fitted into the map. Ribbon models are shown in different colors.

## Discussion

The functional role of PBSs as a major light-harvesting antenna of photosynthesis in cyanobacteria and red algae has been known for decades. Yet, the PBS assembly pathway remained incompletely understood. In this study, we provide insight into the physiological changes that occur in the absence of fully assembled PBS and present molecular-resolution snapshots of cyanobacterial cells to unravel the intermediate steps of PBS assembly under near-native conditions. We focused our efforts on the PBS linker protein ApcE, which modulates the structural organization of the PBS core cylinders^6,9,35^, serves as a terminal emitter in energy transfer pathways^18^, and connects the PBS core with PSII and PSI, as well as the thylakoid membranes^36–38^. We find that, in addition to growth and pigment defects, the removal of ApcE in *Synechococcus sp.* PCC 7002 leads to the formation of micrometer-scale crystalline arrays of PBP rods that emit an unusual fluorescent signal *in vivo*. Together, these data further demonstrate the critical role of ApcE and provide insight into the PBS assembly pathway.

Removal of the ApcE protein from cyanobacteria results in significant phenotype changes. Previous characterizations of ApcE knockouts in cyanobacteria reported assembled PBS peripheral rod cylinders^17^, but did not observe fully assembled PBSs^16,17^. Therefore, removing ApcE prevents the assembly of PBS cores and the complete PBS structure^6,16,17^, which impairs energy transfer to PSI and PSII from PBS in Δ*apcE*. We observed a growth defect in Δ*apcE* (Fig. 1, B and C), a direct consequence of their decreased ability to transfer energy from PBS to PSII and PSI as observed in 77K fluorescence spectra (Fig. 1, H and I). This decrease in growth rate is consistent with previously reported growth rate defects in apcE knockouts of PCC 7002 and PCC 6803^6,14,17,37^. The partial recovery of growth rate on lower agar percentage plates (Fig. 1C) is consistent with other PBS mutants^27^. Recovery of the growth rate to WT levels in Δ*apcE^+^* confirms that the absence of ApcE drives this growth defect.

ApcE removal also affects the pigmentation of cells. There is a decrease in both phycobilin and chlorophyll absorption in Δ*apcE* (Fig. 1D). Phycobilin absorption represents PBPs, while chlorophyll absorption represents PSI and PSII. A decrease in phycobilin absorption was previously reported in *apcE* knockouts of the cyanobacteria *Synechocystis sp*. PCC 6803^17,37^, but *apcE* knockout mutants in PCC 7002 were reported to synthesize a normal amount of PBPs^6^. In low-light and red-light conditions where PBS are unable to harvest light energy, cyanobacteria compensate through increasing expression and rod length of the light-harvesting PBS, resulting in a higher phycobilin/chlorophyll ratio^5,39,40^. Despite a decrease in phycobilin content, we observed an increased phycobilin/chlorophyll ratio in Δ*apcE* absorbance curves. While the decrease in phycobilin absorbance in Δ*apcE* was unexpected, several processes regulate PBS protein content. PBS gene expression is sensitive to the redox state of the cell^41^, and is likely also regulated through light-dependent transcription factors. Furthermore, PBS rods and cores are actively degraded in certain nutrient-limiting conditions^24,42,43^. Δ*apcE* is not starved for nutrients like nitrogen or phosphorus, so it is unlikely that active nblA-dependent PBS degradation is occurring. However, high levels of unassembled PBPs present in Δ*apcE* may be more easily targeted for degradation, or they may inhibit transcription of PBS genes.

Cyanobacteria cells must maintain a proper ratio of energy transfer through PSII and PSI to prevent oxidative stress and to maintain concentrations of ATP and reducing equivalents in the cell^23,44^. PBSs play an important role in this process through modulation of their energy transfer to PSI and PSII through association and dissociation with the photosystems as needed^23^. The Δ*apcE* mutant lacks core-containing PBSs, which favor energy transfer to PSII^45^, but does contain fully assembled CpcL-containing rod-only PBSs that favor energy transfer to PSI^3,46^. Therefore, the cells must compensate for the decreased energy flux through PSII compared to PSI in Δ*apcE*. In the analogous situation where cyanobacteria are illuminated with only PSI-preferred red light, an increase PSII/PSI ratio was observed^47^. Therefore, we investigated changes in chlorophyll and the PSII/PSI ratio in Δ*apcE*. The amount of chlorophyll present in Δ*apcE* was lower than WT (Fig. 1E). Despite the decrease in the amount of chlorophyll in cells, there was an increase in chlorophyll fluorescence at room temperature (Fig. 1F). Chlorophyll fluorescence at room temperature is mostly attributed to PSII^7^, so increased fluorescence typically represents some combination of the following: 1) increased PSII protein levels, 2) decreased PSII photochemistry, and/or 3) decreased non-photochemical quenching of PSII. 77K fluorescence measurements of Δ*apcE* revealed an increase in PSII/PSI ratio, which was also reported in an ApcE knockout strain of PCC 6803^17^ (Fig. 1H). Furthermore, PSII has about 40 chlorophyll molecules per monomer, while PSI has about 100 chlorophyll molecules per monomer; therefore, the increase in PSII/PSI ratio is consistent with a decrease in chlorophyll. The increase in the PSII/PSI ratio is a mechanism to compensate for the decreased transfer of energy from core-containing PBSs to PSII. While it is possible that decreased photochemistry and/or decreased non-photochemical quenching may also contribute to increased chlorophyll fluorescence in Δ*apcE*, we did not characterize these processes.

The chlorophyll fluorescence peak is shifted slightly left at RT in Δ*apcE* (Fig. 1F). This change is likely due to the shoulder of the chlorophyll fluorescence curve at wavelengths corresponding with the extremely fluorescent PBPs in Δ*apcE*. The phycobilin fluorescence peak in Δ*apcE* at RT is also left shifted (Fig. 1G). The phycobilin fluorescence shift is likely driven by several factors, including the absence of the longer wavelength (680 nm emission peak) terminal emitter in ApcE and decreased energy transfer from PC rods (650 nm emission peak) to APC cores (660 nm emission peak) due to impaired PBS assembly. While Δ*apcE* cells exhibit reduced phycobilin absorbance, their phycobilin fluorescence is markedly higher at room temperature (Fig. 1G). The high fluorescence phenotype is characteristic of knockouts of ApcE and ApcA/B^16^. The increased phycobilin fluorescence indicates decreased energy transfer within assembled PBSs and from PBSs to PSI and PSII. In agreement with this conclusion, 77K spectra revealed an increase in unassembled PBPs and free PBSs compared to PSI- or PSII-coupled PBSs in Δ*apcE* (Fig. 1I). Although fully assembled core-containing PBSs were absent, energy transfer from PBSs to PSI and PSII was still detected in the 77K spectra of ΔapcE. This energy transfer is attributed to CpcL-containing rod-only PBSs and/or association of partially assembled PBS core subunits with PSI or PSII. For example, the A0913 protein associates with ApcF, ApcB, and PSII to help link the PBS core to PSII^48^. The 77K fluorescence spectra of Δ*apcE* also confirmed the presence of both PBS rod (PC, 650 nm) and core (APC, 660 nm) proteins (Fig. 1I). Therefore, despite impaired assembly of PBS cores, PBS rod and core proteins are both still present in the cell, consistent with previous findings^17^.

The subcellular spatial organization of Δ*apcE* was observed with fluorescent microscopy of the naturally occurring pigments in cyanobacteria (Fig. 2). Two microscope fluorescent channels were used – the phycobilin-specific RFP channel and the Cy5 channel, which excites phycobilin and the high-energy edge of chlorophyll and collects both chlorophyll and low-energy phycobilin fluorescence. The presence of fluorescent puncta in both the RFP and Cy5 channels that have correlated values and positions supports the existence of highly fluorescent PBP aggregates in Δ*apcE* that are not present in WT or Δ*apcE^+^.* These PBP aggregates tended to be pushed towards the edge of the long axis of Δ*apcE* cells. In bacteria, the nucleoid acts as a diffusion barrier for large structures, and exclusion by the nucleoid pushes large subcellular structures such as inclusion bodies or carboxysomes away from the center of the cell in the absence of active positioning systems^49,50^. When the carboxysome-positioning system is knocked out, the localization of the carboxysomes at cell poles is similar to the positioning of PBP aggregates in Δ*apcE*^49^. Therefore, the subcellular position of PBP aggregates near the poles of cells is likely due to exclusion by the nucleoid and not an active positioning system. It is unlikely that the RNA of PBS components is specifically localized to cell poles for translation, as the RNA for PC was shown to be significantly more cytoplasmic and less associated with thylakoid membranes and “biogenesis centers” than RNA for components of PSI and PSII^51^.

Time-lapse microscopy revealed that fluorescent PBP aggregates persist over multiple cell divisions in a stable subcellular position in Δ*apcE* (Fig. 3). While PBP aggregates were frequently inherited by only one daughter cell after division, occasionally a PBP aggregate was split between daughter cells. If an *apcE* cell did not inherit at the PBP aggregate during cell division, a PBP aggregate formed within a few hours.

Development of an inducible CRISPRi knockdown of ApcE allowed additional analysis of PBP aggregates through time-lapse microscopy. PBP aggregates form about 10 hours after ApcE knockdown is induced in time-lapse microscopy and persist in stable subcellular positions like those in Δ*apcE*. When induction of ApcE knockdown in the CRISPRi strain ceased, PBP aggregate formation stopped, and fluorescence of existing PBP aggregates faded over time. Therefore, PBP aggregate formation is reversible, since fading fluorescence of PBP aggregates during cell growth represents the disassembly of the aggregate.

Cryo-CLEM and subsequent cryo-ET of Δ*apcE* revealed that these highly fluorescent PBP aggregates are large crystalline structures (Fig. 4). A 3D reconstruction was created through sub-tomogram averaging of the crystalline PBP aggregates (Fig. 5). In addition to similar molecule sizes between the crystalline structure observed in Δ*apcE* and PBS cylinders, there was strong agreement between our 3D reconstruction and the cryo-EM structure of peripheral PBS rods from PCC 7002^9^. However, instead of cylinder lengths between 2 and 6 hexamer discs like those observed in assembled PBSs, the cylinders observed in the Δ*apcE* cells have the length of dozens of hexamer discs. The long cylinders in Δ*apcE* pack in parallel, which is similar to cylinder packing observed in the rods of fully assembled PBSs stacked on thylakoid membranes in electron microscopy of cyanobacteria^52^. A similar rod packing was also observed in purified PC hexamer structures from the cyanobacteria species Thermoleptolyngbya sp. O-77^53^, Thermosynechococcus vulcanus^54^, and Thermosynechococcus elongatus^55^. Parallel packing of PBS cylinders may facilitate transfer of light energy between the stacked PBSs *in vivo* in assembled PBSs^9^, and likely contributes to the extremely high fluorescence signal of the PBP aggregates in Δ*apcE*.

Interestingly, fluorescent puncta and PBP cylinder aggregates similar to those seen in Δ*apcE* have been purified from a *Synechocystis sp.* BO 8402, a nitrogen-fixing cyanobacterial species lacking APC and assembled PBS cores^56,57^. While we were unable to determine if the crystalline protein aggregate in Δ*apcE* contains rod linker proteins, rod linker proteins were identified in the PBP cylinder aggregates of BO 8402^56^. Since similar crystalline PBP aggregates are observed in different strains of cyanobacteria where PBS core assembly is prevented through the absence of different PBS core components, we assume the crystalline PBP structures represent the accumulation of a PBS assembly intermediate. When ApcE expression is recovered in the CRISPRi knockdown strain, PBP aggregate disassembly likely represents the incorporation of PBS assembly intermediate into fully assembled PBSs. Furthermore, an increase in first quartile (non-puncta) fluorescence in the RFP channel of Δ*apcE* observed in light microscopy could represent increased concentrations of early PBS assembly intermediates like free PBP monomers, trimers, and hexamers that are soluble in the cytoplasm (fig. S2) These data, along with the fluorescent and structural characteristics of the crystalline aggregates, demonstrate that assembled peripheral rods are an intermediate in PBS assembly.

Protein aggregation is a frequent event in biological systems. Common examples of protein aggregates include amorphous aggregations of unfolded proteins formed in stress response, called inclusion bodies, and paracrystalline aggregations of misfolded proteins into amyloid fibrils^58^. In rare instances, properly folded proteins aggregate in a crystalline manner like the structures observed in Δ*apcE*. Crystalline protein aggregates in bacteria have been found to function in storage, encapsulation, or compartmentalization^59^. All cyanobacteria possess protein-bound organelles called carboxysomes, which have small heterogeneous two-dimensional crystalline protein lattices forming the icosahedral shell that compartmentalizes carbon fixation enzymes^59–61^. Many archaea and bacteria species, including some cyanobacteria, have S-layers, which are additional examples of a two-dimensional crystalline protein lattice that encapsulates the entire cell and performs multiple functions^59,62,63^. PBSs are not involved in encapsulation or compartmentalization, but instead are involved in nutrient storage^24,64^. Therefore, the crystalline structures observed in Δ*apcE* may play a role in nutrient storage in addition to PBS assembly.

The ability to produce such a large protein aggregate of a PBS intermediate without triggering a degradation pathway or activating PBS gene repression is perhaps symbolic of the existence of a role for PBP aggregation in the PBS assembly pathway. It is known that PBS cores and PBS rods can assemble independently^65^, but the final steps of the PBS assembly process have not been well characterized. While core-containing PBSs and CpcL-containing rod-only PBSs both contain assembled peripheral rods, they favor association with different photosystems. Furthermore, PBS cores will associate with and transfer energy to PSII in the absence of peripheral rods^17,65,66^. We propose a model of PBS assembly where CpcL linker protein or assembled PBS cores associated with newly assembled PSI or PSII, respectively, pull assembled peripheral rods from a pool maintained in the biogenesis centers (Fig. 6). The impaired the assembly of PBS cores in the Δ*apcE* mutant reduces flow out of the peripheral rod pool, increasing the concentration of free peripheral rods and facilitating the formation of the large crystalline PBP structure. In this model of PBS assembly, the flux of light energy through PSI and PSII in the long term could be regulated through CpcL and PBS core protein expression, which compete for the same pool of PBS peripheral rods^67^. CpcL is not part of the PBS core operon or the PBS peripheral rod protein operons, and the differential regulation of CpcL and other PBS genes provides support for this _model26,68,69._

**Figure 6.**
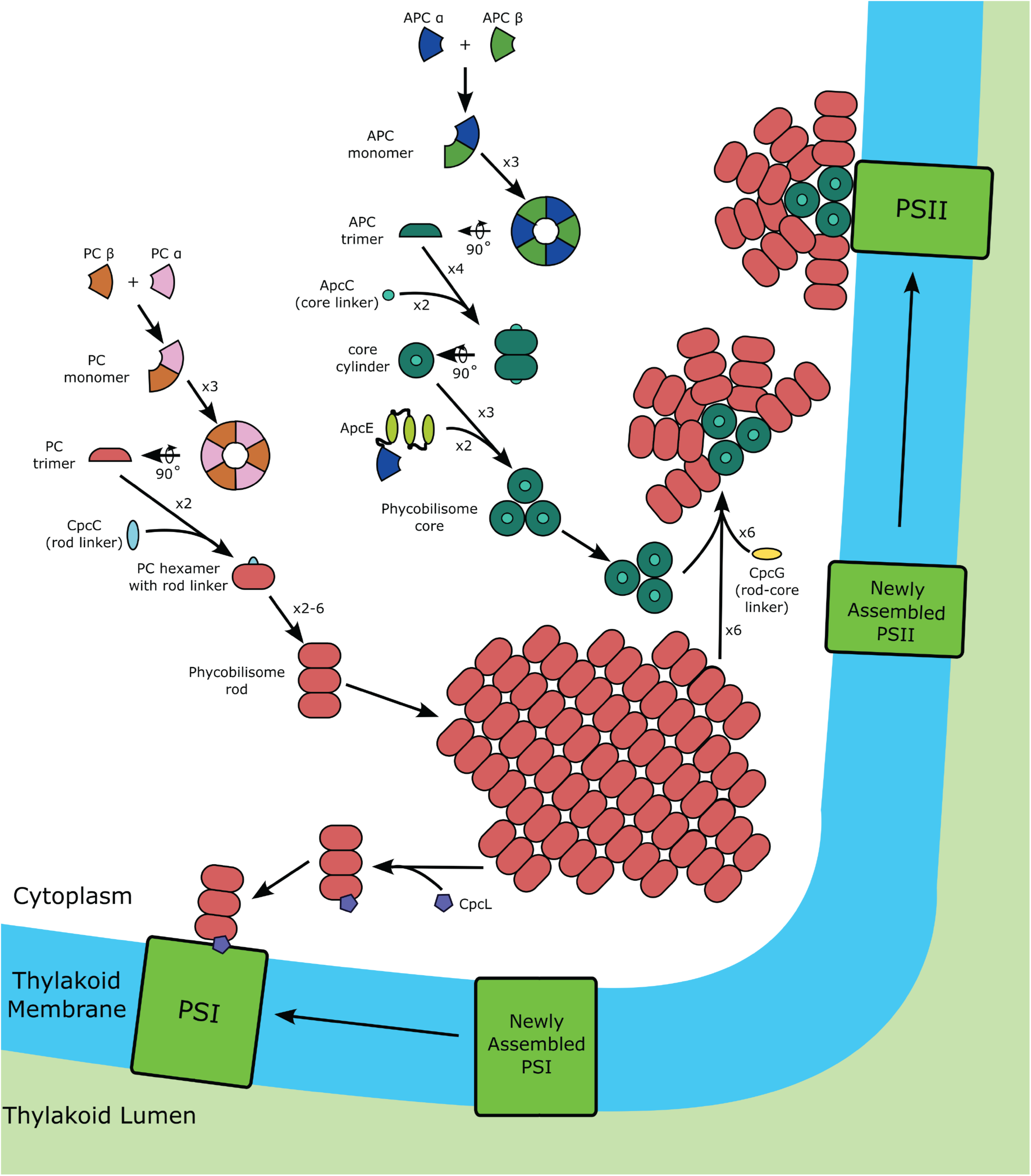
Model of PBS assembly with peripheral rod pool intermediate. A model of PBS assembly in PCC 7002. PBS peripheral rod and core assembly occurs in the cytoplasm and assembled PBSs associate with PSI and PSII, large protein complexes in the thylakoid membrane. CpcG-containing PBS cores or CpcL compete for a small pool of PBS peripheral rods. CpcL rod-only PBS typically associate with PSI, while core-containing PBS typically associate with PSII.

In summary, our study highlights the important role of ApcE in coordinating the assembly of PBSs, which support cyanobacterial growth through their role in energy capture and transfer. Physiological and spectroscopic assays of Δ*apcE* demonstrated that the absence of ApcE disrupts PBS core assembly and provided insight into how cyanobacteria regulate the distribution of excitation energy between PSI and PSII. Furthermore, a combination of light and electron microscopy revealed the accumulation of crystalline PBP arrays, a stalled intermediate in the PBS assembly pathway. The structures observed in Δ*apcE* provide a striking example of crystalline protein aggregation. Together, these findings advance our understanding of PBS biogenesis and function, supporting a new model of PBS assembly, and have broader implications for improving energy transfer efficiency in other organisms and bioengineering applications.

## Materials and Methods

### Creation of Δ*apcE* and Δa*pcE*+ strains

The *apcE* knockout plasmid contains gentamycin resistance between *apcE* up and down homology arms (600 bp each) on a pUC19 backbone. The *apcE* complement plasmid contains spectinomycin resistance and *apcE* behind pK2^70^, a strong constitutive promoter, between neutral site 1 (NS1)^71^ homology arms on a pUC19 backbone. For the *apcE* knockout and complement plasmids, homology arms, *apcE,* and pK2 were amplified from PCC 7002. Plasmids were assembled using Gibson Assembly^72^. The Gibson reactions were transformed into DH5α *E. coli* and minipreps of liquid cultures started from single colonies were performed to collect plasmid. Plasmid was transformed into PCC 7002^73^ and colonies containing the desired insert were serially passaged in the presence of antibiotic until segregated. The primers used to confirm segregation of *apcE* and NSI loci are listed in table S1.

### Creation of CRISPRi strains

A previously developed IPTG-inducible sgRNA expression plasmid^74^ was modified using Gibson assembly and cloned into DH5α *E. coli*. Homology arms for insertion downstream of the *glpK* genomic locus^71^ were amplified directly from PCC 7002 cells. The *apcE* sgRNA spacer (5’-GAACGGTTTGATATAACTGG-3’) was designed to bind the coding strand near the beginning of the *apcE* coding sequence. The non-targeting (NT) spacer (5’- TCTTTGGGTCGCCCGTCGAC-3’) harbors no significant homology in the PCC 7002 genome. Spacers were inserted using Golden Gate assembly^75^. Each sgRNA expression cassette was transformed into PCC 7002^73^ containing dCas9 under Tet inducible control in the *acsA* locus^76^ and serially passaged in the presence of antibiotic until segregated.

### Growth conditions

Liquid cultures were used for growth curves, absorbance scans, and fluorescence measurements. Liquid 50 mL cultures of PCC 7002 strains were grown in shaking flasks containing A+ media^77^ in air at 37°C with a light intensity of 185 µmol photons • m^−2^ • s^−1^. PCC 7002 strains were grown on A+ plates with 1% or 0.5% agar percentages in air at 37°C with a light intensity of 115 µmol photons • m^−2^ • s^−1^. To induce expression of dCas9, 0.5 µg/mL of anhydrotetracycline (aTC) was included in A+ plates or liquid cultures. To induce expression of sgRNA, 5 mM of Isopropyl β- d-1-thiogalactopyranoside (IPTG) was included in A+ plates or liquid cultures.

### Growth curve measurement

The growth of PCC 7002 was measured in liquid cultures grown as described. These cultures were inoculated in triplicate with 1 mL of PCC 7002 pre-culture diluted to 0.271 OD_730_. The precultures were started from PCC 7002 cells scraped from plates and grown in the same conditions as the growth curve cultures, including addition of aTC and/or IPTG. During the growth curve, time points were taken every 3 hours for 48 hours, every 6 hours for the next 24 hours, and every 8 hours for the remaining 48 hours of the experiment. At each time point, 200 µL was removed from each culture and the OD_730_ was measured in a 96-well plate on a Tecan Spark multimode microplate reader.

### Spot plates

Plates with and without each inducer and at 0.5 and 1% agar were spotted with uninduced strains in triplicate. Liquid cultures of each uninduced strain of PCC 7002 were diluted to 0.1 OD_730_ and six 1:10 serial dilutions were performed. Each row on a spot plate was one set of serial dilutions. Seven µL of the serial dilutions was used for each spot. Images were taken 3 days after spotting the plates.

### Room temperature absorbance and fluorescent spectra measurements

The cultures used for room temperature absorbance and fluorescence spectra were inoculated with cells from liquid cultures containing indicated inducers. The cultures were grown in liquid medium with indicated inducers to 0.3-0.5 OD_730_ in triplicate. The cultures were diluted to 0.183 OD_730_ and 200 µL of each culture was loaded onto a 96-well plate. Room temperature absorbance and fluorescence were measured on a Tecan Spark multimode microplate reader. The absorbance was measured from 400-750 nm with a step size of 2 nm and a bandwidth of 3.5 nm. Phycobilin fluorescence emission was measured from 600-800 nm with a bandwidth of 20 nm and a step size of 2 nm using an excitation wavelength of 580 nm with a bandwidth of 20 nm. Chlorophyll fluorescence emission was measured from 600-770 nm with a 20 nm bandwidth and a step size of 2 nm using an excitation wavelength of 440 nm with a bandwidth of 20 nm.

### Chlorophyll quantification

The cultures used for chlorophyll measurements were inoculated from pre-cultures grown in liquid cultures containing indicated inducers. Liquid cultures were grown with indicated inducers to OD 0.3-0.5 in triplicate. Chlorophyll was methanol extracted from 1 mL of culture diluted to 0.233 OD_730_ as described in Porra et al.^78^ Absorbance at 665 nm was measured and the chlorophyll content was calculated with equation 1.

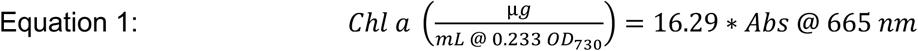

### Fluorescence measurements at 77 K

The cultures used for 77K fluorescence measurements were inoculated with cells from liquid cultures containing indicated inducers. The cultures were grown in liquid medium with indicated inducers to OD 0.3-0.5 in triplicate. OD_730_ was measured and 1 mL of each culture at 0.233 OD_730_ was made by diluting the original culture and was used for chlorophyll quantification. Each original culture was diluted to a chlorophyll concentration of 2 µg/mL in 10 mLs. The normalized chlorophyll cultures were pelleted at 4300 x g for 10 min at room temperature and the supernatant was removed. The pellet was resuspended in 1 mL A+ and 600 µL was loaded into a medium-walled NMR tube and frozen in liquid N_2_ for 77 K fluorescence measurements. 77K fluorescence was measured using a Fluorolog-3 spectrofluorometer (Horiba Jobin Yvon) with a FL-1013 attachment to allow for measurement of samples submerged in liquid N_2_. Phycobilin fluorescence emission at 77 K was measured from 600-800 nm with a 5 nm slit and a step size of 1 nm using an excitation wavelength of 580 nm with a 5 nm slit. Chlorophyll fluorescence emission at 77 K was measured from 600-770 nm with a 5 nm slit and a step size of 1 nm using an excitation wavelength of 440 nm with a 5 nm slit.

### Widefield time-lapse microscopy

Time-lapse widefield microscopy was performed using the microscope and software described in Moore *et al.* and Hill *et al.*^27,79^. Briefly, images were taken using a customized Nikon TiE inverted widefield microscope with a near-infrared-based Perfect Focus System, a custom Okolab environmental enclosure for temperature and CO_2_ control, an individually controllable RGB LED light source for transillumination (Lida Light Engine, Lumencor), a high-speed light source with customized filter sets for imaging (Spectra X Light Engine, Lumencor) and a synchronized and hardware-triggered motorized shutter for light control. The microscope was controlled using NIS Elements AR software (version 5.11.00 64-bits) with the Jobs acquisition upgrade. Images were acquired on an ORCA Flash4.0 V2+ digital sCMOS camera (Hamamatsu) using a Nikon CFI60 Plan Apochromat Lamda ×100/1.45 NA oil-immersion objective.

For time-lapse microscopy, 2 µL of cells in exponential phase were spotted onto a 1% or 0.5% agarose A+ pad, which was placed onto a 35 mm glass-bottom dish (Ibidi) which was wrapped with parafilm to prevent water loss from the agarose pad. The agarose pad contained aTC and IPTG when indicated. The cyanobacteria were grown in air at 37°C during time-lapse microscopy. Fluorescent and brightfield images were taken every 20 min. Chlorophyll and phycobilin fluorescence was imaged using a 640 nm LED light source (Spectra X), and emission wavelengths were collected using a standard Cy5 filter (Nikon). Phycobilin fluorescence was imaged using a 555 nm LED light source (Spectra X), and emission wavelengths were collected using an RFP filter (571-628 nm) (Nikon). For brightfield images, transmission of RGB LED light through the samples was collected. Cells were constantly illuminated with red light except during imaging.

### Cryo-ET sample preparation

Frozen, hydrated specimens were prepared as described previously^74^. Briefly, Quantifoil R2/2 mesh 200 holey carbon grids were glow-discharged for 45 seconds at 15 mA to increase hydrophilicity. A 4 μL aliquot of bacterial cell suspension was applied to the carbon side of the grid, followed by incubation at 80% humidity and 22°C in the Leica EM GP2 plunge freezer chamber. Excess liquid was removed by one-sided blotting. Blotting conditions were varied to obtain optimal ice thickness and cell count. Generally, a 1 to 2 minute incubation time was followed by a 2-3 second blotting time using filter paper (Whatman) from the opposite side of the grid and immediate plunging into liquid ethane cooled by liquid nitrogen. The vitrified samples were subsequently transferred to storage boxes and maintained in liquid nitrogen until further processing.

### Cryo-fluorescence light microscopy

To verify the cell quantity on the EM grid before FIB-SEM milling, vitrified grids were imaged using a THUNDER Imager EM Cryo CLEM system (Leica) equipped with a cryo-stage maintained at −196°C to prevent sample devitrification. Widefield fluorescence overview montages were acquired to identify grid squares containing cells of interest. A wavelength corresponding to the red fluorescence channel (596\615 excitation\emission) was used to observe chlorophyll fluorescence emitted from cyanobacterial cells, confirm uniform sample coverage, and validate the presence of the typical fluorescence puncta for the Δ*apcE* strain on the pole of the cell.

### Cryo-focused ion beam milling

Clipped grids were transferred to an Aquilos dual-beam FIB-SEM microscope (Thermo Fisher Scientific) equipped with a cryo-transfer system and a 360° rotatable cryo-stage. Throughout the FIB operation, samples were maintained in high vacuum (<4 × 10⁻⁷ mbar) at constant liquid nitrogen temperature. Initial low-magnification SEM imaging (2kV, 25 pA) was performed to locate the previously identified regions of interest based on grid square patterns and correlated light microscopy data. Before milling, samples were sputter-coated with platinum (1 kV, 20 mA, 25 seconds at 0.10 mbar) to improve conductivity and reduce charging artifacts. Additional organometallic platinum was deposited using the gas injection system (GIS) operated at 28°C with a 7 mm stage working distance and 90-second gas injection time to provide protection during the milling process. Lamella preparation was performed using a series of milling steps with sequentially decreasing ion beam currents. Initial rough milling was conducted at 1 nA to create trenches on both sides of the region of interest, followed by intermediate milling at 300 pA, fine at 100 pA, and finer at 100 pA. Final polishing was performed at 50 pA and 10 pA to achieve a final lamella thickness of approximately 150-200 nm suitable for transmission electron microscopy. The fabricated lamellae were positioned perpendicular to the grid plane at a shallow angle of 12° relative to the grid surface to maximize the observable area within the bacterial cells.

### Cryo-ET data collection

Lamellae were imaged using a 300 kV Titan Krios G3i transmission electron microscope equipped with a Selectris energy filter and Falcon 4i electron detector (Thermo Fisher Scientific). Tilt series were collected using Tomography 5 software with a dose-symmetric tilt scheme ranging ±55° with 3° increments around the pre-tilt angle defined by the milling angle, resulting in a total of 57-60 projections per tilt series. Images were acquired at a nominal magnification of 81,000×, corresponding to a pixel size of 1.76 Å at the specimen level. The cumulative electron dose for the tilt series was kept at approximately 110 e⁻/Å². Defocus values ranged from −2 to −8 μm, and energy filtering was performed with a 10eV slit width.

### Cryo-ET data processing and 3D reconstruction

Initial data processing was performed in Inspec3D (Thermo Fisher Scientific) and EMAN2 (version 2.99.66)^80^ to assess data quality and initial feedback on structure determination. For this step, motion correction of movies was performed in IMOD-align frames. Inspect3D allowed a quick review of tilt series quality and tomogram reconstruction to identify the presence of target crystals/particles and evaluate thickness. The best tilt series were selected. Further processing and subtomogram averaging were carried out using RELION (version 4.0.2)^34^. Movies were motion corrected with the RELION motion correction implementation. Tilt series alignment was done in IMOD (version 5.1)^32^ using Etomo with patch tracking and manual supervision/optimization of parameters/steps for each tilt series. CTF estimation was done first in IMOD-Ctfploter^81^, and this was used to optimize the search parameters in CTFFIND4^82^. Pseudo-subtomos of a first set of 312 particles, manually picked. A full set of 32907 particles was picked using crYOLO (version 1.9.9)^83^ and extracted with a cropped box size of 192 pixels and 2x binned. Three initial models were generated with a 500Å mask. Best initial model and 4145 particles were used for refine3D with a 500Å mask, C1 symmetry initial low pass 60Å. Map from refine3D was used for final post-processing, which resulted in a ∼19 Å resolution FSC. A lattice correlation map was generated in RELION with the tomo find lattice module, using the tomogram generated in IMOD, and the distance was measured from the tomogram, aligning columns perpendicular and measuring from the center to the center of columns. All cryo-ET data processing and analysis software was compiled and supported by the SBGrid Consortium^84^. The data acquisition and image processing parameters are summarized in table S2. An overview of cryo-ET data collection and image processing statistics was provided in Fig. S3.

### Model building, refinement, and validation

The cryo-EM structure of *Synechococcus sp.* PCC 7002 PBS (PDB ID: 7EXT) was used as an initial reference for model fitting and refinement. The hexameric models were extracted from the complex structure and manually fitted into the density map using both Chimera (version 1.19)^85^ and ChimeraX (version 1.9)^86^. After model fitting, the model was refined against the cryo-ET map using phenix.real_space_refine tool in the PHENIX software package (version 1.21.2)^87^. The PHENIX model-map correlation coefficient validated the quality of the model fitting. The structural figures were prepared in Chimera using the *in situ* cryo-ET map and hexameric PBS peripheral rod models.

## Acknowledgments

The authors would like to dedicate this paper to the memory of Prof. Jeffrey C. Cameron, who provided funding and served as a mentor throughout most of this work. We thank the National Network for Cryo-Electron Tomography, an NIH-funded consortium, for electron cryo-tomography training and assistance with sample preparation and data collection; the members of the Cameron and Aydin labs for helpful discussions and critically reading and editing the manuscript. We also thank Ash Weier and Zoltan Metlagel for assistance with cryo-ET data collection and processing, and Evan Johnson and Dr. Clair Huffine for their support with time-lapse imaging of Δ*apcE*.

## Authors contributions

K.D., A.A., E.K., J.C.C., and H.A. designed the project. E.K. and A.H. designed and produced strains. K.D., A.A., E.K., A.G.I., A.H., and L.G. performed experiments and collected experimental data. A.A, A.G.I., F.J. A-R., A.C., J.H., and J.S. carried out electron cryo-tomography imaging and image analysis. K.D., A.A., E.K., A.G.I., J.C.C., and H.A. analyzed the data and discussed the results. J.C.C. and H.A. supervised the project. K.D., A.G.I., and H.A. wrote the manuscript with the input from all authors.

## Supplementary Figure Legends

**Figure S1.**
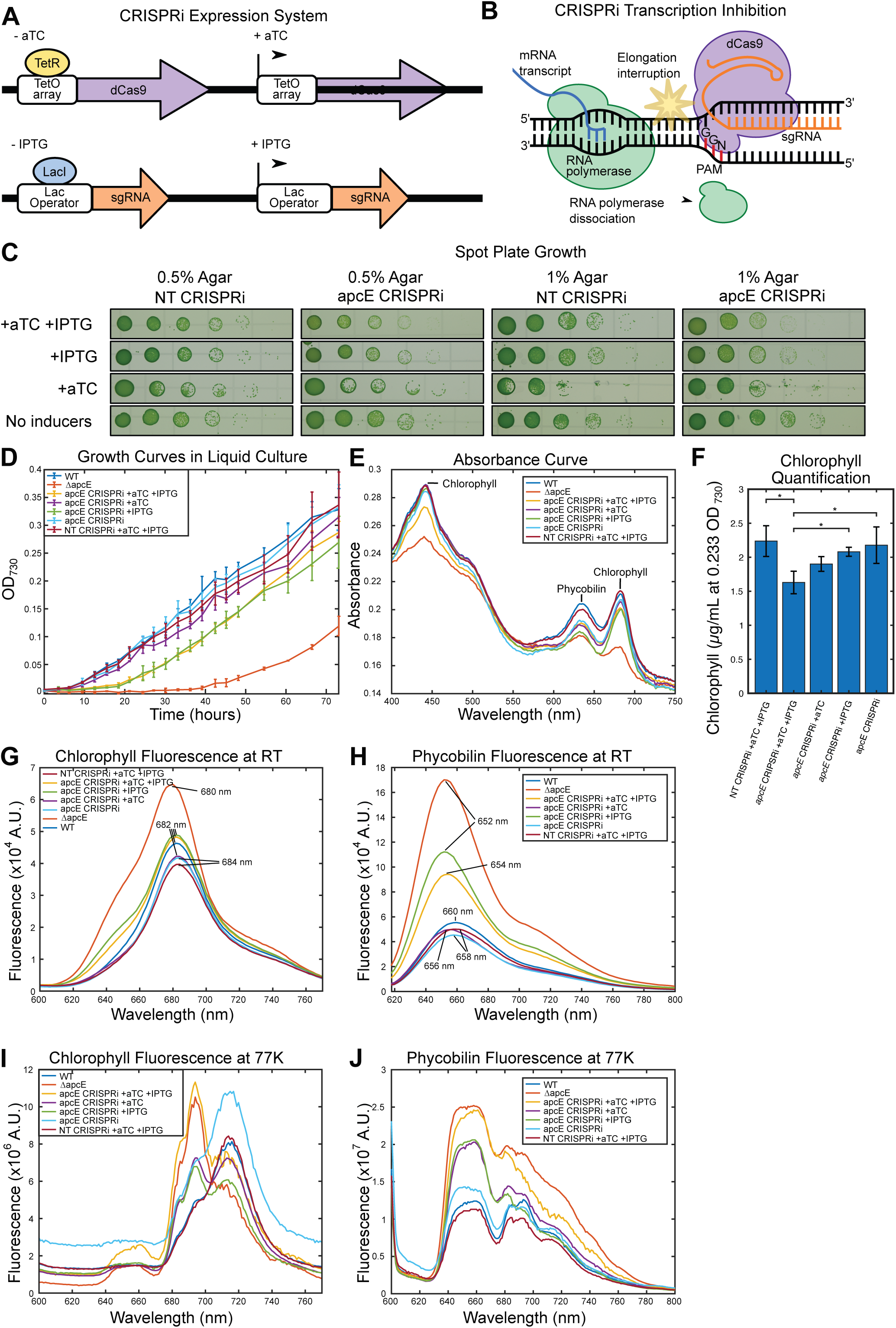
***apcE* knockdown has growth and pigment defects.** (**A**) The top shows induction of dCas9 through addition of aTC. In the absence of aTC, TetR is bound to a TetO array, preventing transcription of dCas9. When aTC is present, TetR dissociates from the TetO array, allowing transcription of dCa9. The bottom shows induction of an sgRNA with IPTG. In the absence of IPTG, LacI binds to the lac operator, preventing transcription of the sgRNA. When IPTG is added, LacI dissociates from the lac operator allowing transcription of the sgRNA. (**B**) A schematic of repression of a target gene by CRISPRi. DNA is shown in black, with individual bases represented by vertical black lines. On the left, RNA polymerase transcribes a gene. The dCas9-sgRNA is bound to the DNA through base pairing on the left. The protospacer adjacent motif (PAM) sequence of 5’-NGG-3’ is marked with red DNA bases. The dCas9-sgRNA bound to the DNA sterically interrupts the elongation of mRNA by RNA polymerase, resulting in the dissociation of RNA polymerase from the DNA. (**C**) The growth of WT, *ΔapcE*, NT CRISPRi with aTC and IPTG, and apcE CRISPRi with and without aTC and IPTG was recorded. OD_730_, a measure of cell density, was collected over 72 hours from cultures inoculated with 1 mL of culture diluted to the same OD_730_. Time is on the x-axis and OD_730_, a measure of cell number, is on the y-axis. The average OD_730_ of three replicates of each strain is displayed. The error bars represent one standard deviation. (**D**) The growth of NT CRISPRi (first and third columns) and *apcE* CRIPSRi (second and fourth columns) at 0.5 % (first and second columns) and 1 % (third and fourth columns) agar plates with and without inducers was compared. The top row has aTC and IPTG present, the second row has only IPTG present, the third row has only aTC, and the bottom row does not have either inducer. Uninduced cultures of NT CRISPRi and *apcE* CRISPRi were diluted to the same OD_730_ and serial 1:10 dilutions were performed for each spot from left to right. A representative image from triplicate spot plates is displayed. (**E**) Absorbance spectra of WT, *ΔapcE*, NT CRISPRi with aTC and IPTG, and apcE CRISPRi with and without aTC and IPTG were collected. All samples were diluted to the same OD_730_ to normalize the cell number and the absorbance spectra were measured. Wavelength is on the x-axis, while absorbance is on the y-axis. Each line is an average of three replicates. (**F**) The amount of chlorophyll was quantified from NT CRISPRi +aTC +IPTG and *apcE* CRISPRi with and without each inducer. Cells from a liquid culture were diluted down to the same OD_730_, chlorophyll was methanol extracted, and the absorbance at 665 nm was measured and used to calculate the amount of chlorophyll per OD_730_. Error bars represent one standard deviation. (**G**) Chlorophyll fluorescence emission spectra of WT, *ΔapcE*, NT CRISPRi with aTC and IPTG, and apcE CRISPRi with and without aTC and IPTG were collected at room temperature. Each sample was diluted down to the same OD_730_ to normalize the cell number. The excitation wavelength was 440 nm. The emission wavelength is on the x-axis and fluorescence is on the y-axis. Each line represents the mean of three samples. (**H**) Phycobilin fluorescence emission spectra of WT, *ΔapcE*, NT CRISPRi with aTC and IPTG, and apcE CRISPRi with and without aTC and IPTG were collected at room temperature. Each sample was diluted down to the same OD_730_ to normalize the cell number. The excitation wavelength was 580 nm. The emission wavelength is on the x-axis and fluorescence is on the y-axis. Each line represents the mean of three samples. (**I**) Chlorophyll fluorescence emission spectra of WT, *ΔapcE*, NT CRISPRi with aTC and IPTG, and apcE CRISPRi with and without aTC and IPTG were collected at 77 K. Each sample culture was normalized to the same chlorophyll concentration. The excitation wavelength was 440 nm. The emission wavelength is on the x-axis and fluorescence is on the y-axis. Each line is a representative emission spectrum selected from three samples of each strain. PSII emission maxima are 685 and 695 nm, while the PSI emission maximum is at 720 nm. (**J**) Phycobilin fluorescence emission spectra of WT, *ΔapcE*, NT CRISPRi with aTC and IPTG, and apcE CRISPRi with and without aTC and IPTG were collected at 77 K. Each sample culture was concentrated to the same chlorophyll concentration. The excitation wavelength was 580 nm. The emission wavelength is on the x-axis and fluorescence is on the y-axis. Each line is a representative emission spectrum selected from three samples of each strain. Free PBS emission maxima are from 640-660 nm, PSII emission maxima are 685 and 695 nm, and PSI emission maximum is at 720 nm.

**Figure S2.**
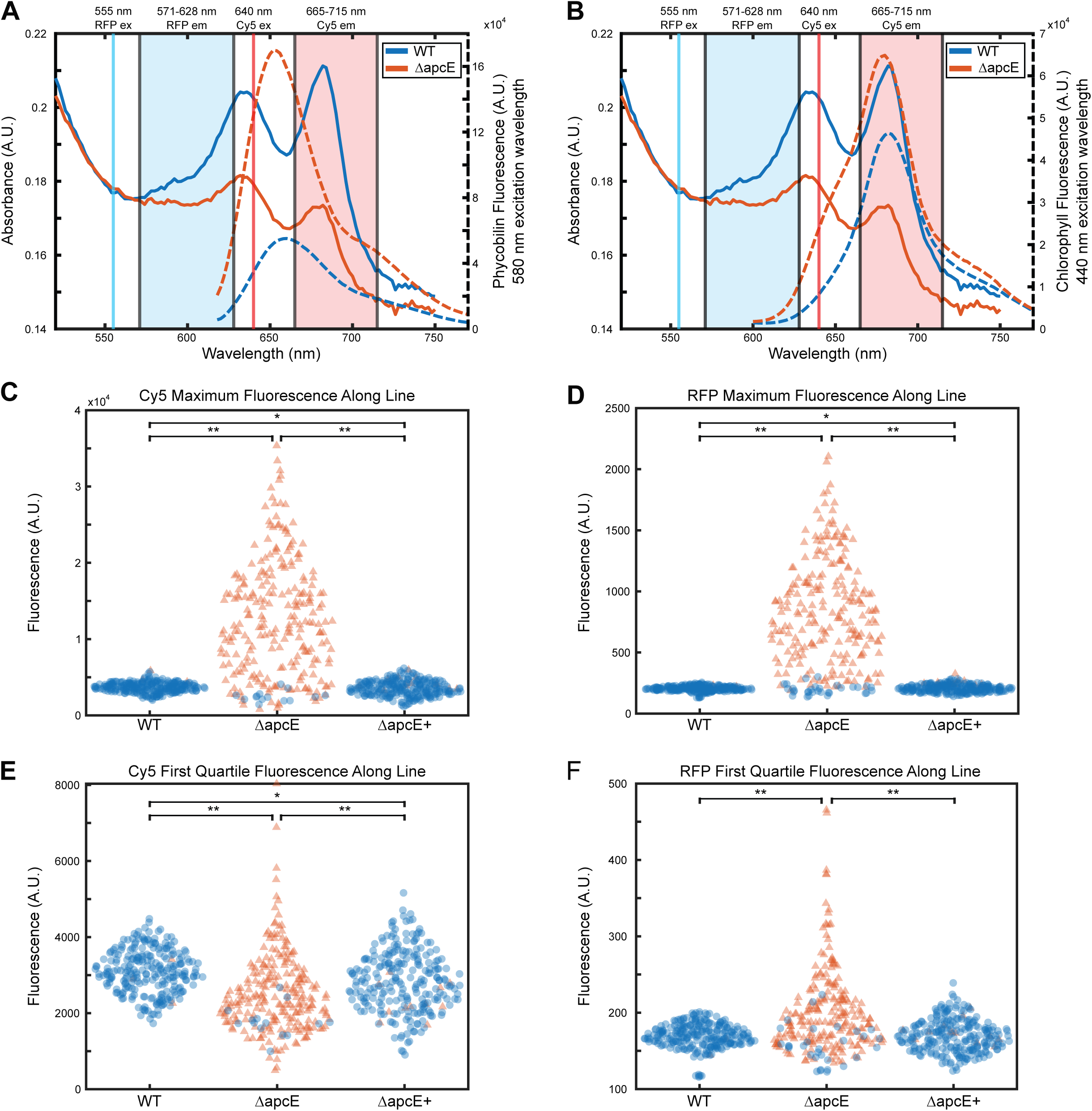
Supplemental fluorescence microscopy of Δ*apcE*. The excitation and emission wavelengths for the Cy5 (red) and RFP (cyan) channels overlaid with the absorbance curve (solid line) of Δ*apcE* (red) and WT (blue) and room temperature (**A**) phycobilin or (**B**) chlorophyll fluorescence (dotted line) of Δ*apcE* (red) and WT (blue). (**C** to **F**) The distribution of maximum (**C** and **D**) or first quartile (**E** and **F**) fluorescence values of a pole-to-pole line in the Cy5 (**C** and **E**) or RFP (**D** and **F**) channels. The distributions of WT, Δ*apcE*, and Δ*apcE+* are shown the left, middle, and right of each panel, respectively. The fluorescence values from 250 cells per strain are plotted. Red triangles represent cells with puncta values greater than 3 times the standard deviation above the WT average in that respective channel. All other points are shown as blue circles. The asterisk (*) symbol indicates p<0.5 and ** indicates p<1*10^-10^.

**Figure S3.**
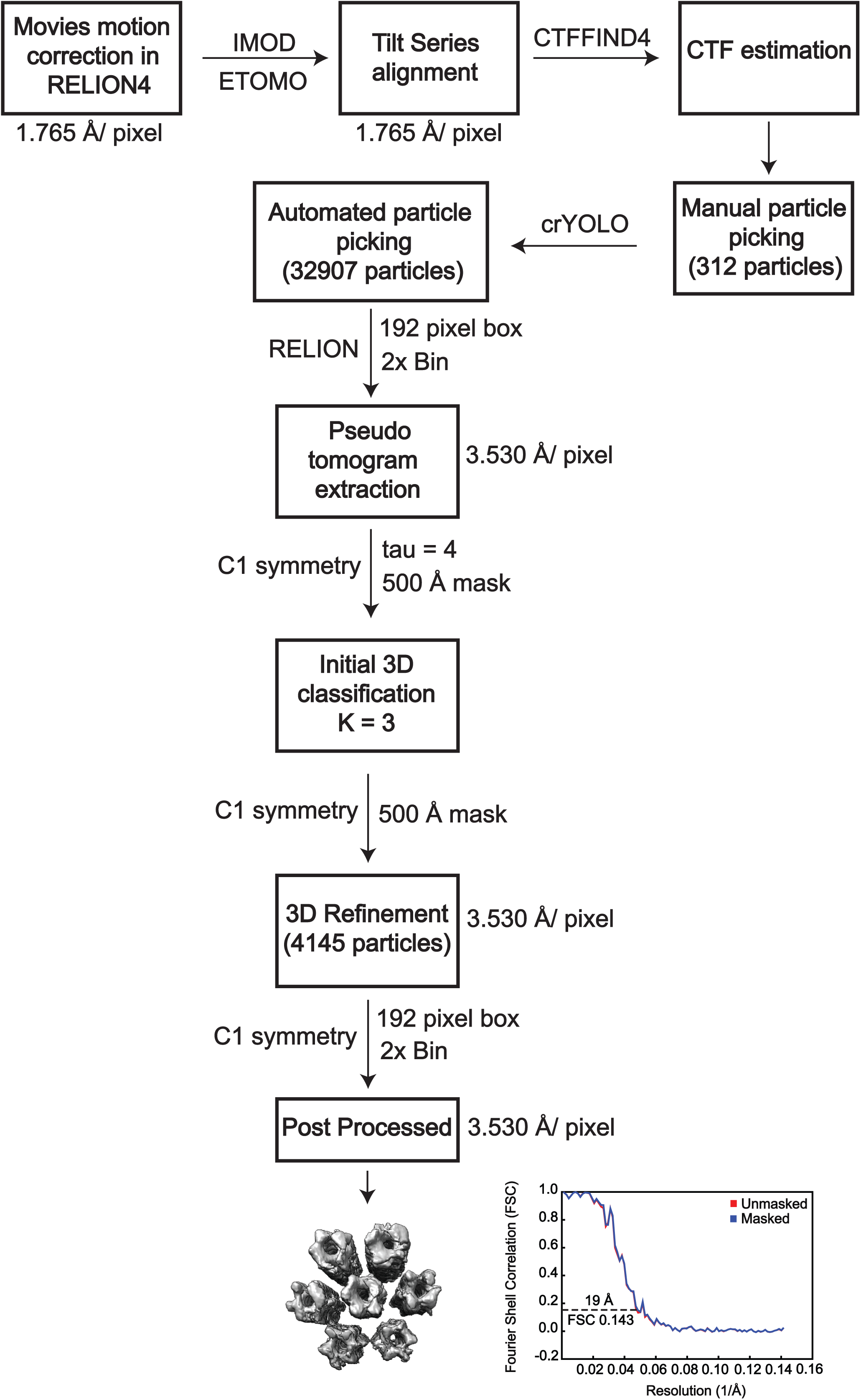
Schematic overview of the subtomogram averaging workflow. *In situ* visualization of crystalline arrays in ΔapcE cells. 3D tomograms were reconstructed and subvolumes were picked in ΔapcE cells. 3D classification and refinement were performed to align and average the subtomogram for the PBS rod bundle. Details of cryo-ET data collection, image processing, and subtomogram averaging are described in the methods section.

**Table S1.**
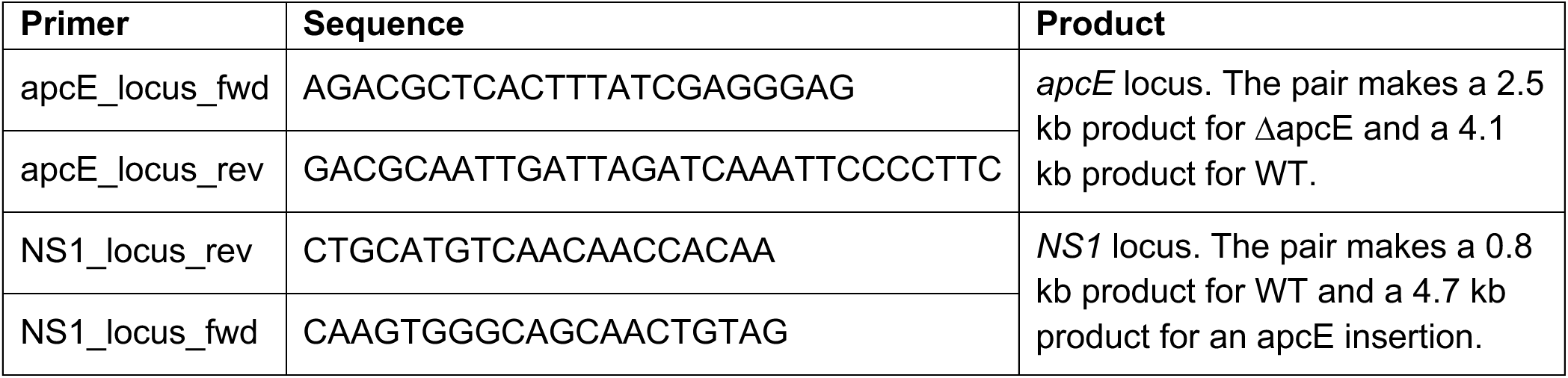
List of cloning primers.

**Table S2.** Cryo-ET data collection and processing statistics for Phycobilisome rod array.

|  |  |
| --- | --- |
|  | **Phycobilisome rod array in *Synechococcus* sp.**<br>**PCC7002**<br>**EMD-73555** |
| **Data Collection and Processing** |  |
| Microscope | FEI Titan Krios |
| Energy Filter / Camera | Selectris / Falcon 4 |
| Magnification | 81,000x |
| Voltage (kV) | 300 |
| Electron exposure (e<sup>-</sup>/Å<sup>2</sup>) | ~110 |
| Defocus (μm) | -2 to -8 |
| Pixel size (Å) | 1.765 |
| Symmetry imposed | C1 |
| Initial segment images (no.) | 32907 |
| Final segment images (no.) | 4145 |
| Map resolution (Å) | 19.34 |
| FSC threshold | 0.143 |

## Supplemental Movie Legends

**Supplemental Movie 1. Δ*apcE* time-lapse.**

Time-lapse fluorescence microscopy was performed to characterize the dynamics of the highly fluorescent puncta. A single colony of Δ*apcE* was imaged every 20 minutes over 94 hours and 20 minutes. The movie displays a merged image of transmitted light (black and white) and autofluorescence in the Cy5 channel (scale in Fig 2A) at every time point. The time point is displayed on the top left and a 2 µm scale bar is displayed in the top right. A green arrow tracks a single punctum throughout the entire time-lapse.

**Supplemental Movie 2. Δ*apcE* time-lapse with cell death and puncta dissipation.**

Time-lapse fluorescence microscopy was performed to characterize the dynamics of the highly fluorescent puncta. A single colony of Δ*apcE* was imaged every 20 minutes over 94 hours and 20 minutes. The movie displays a merged image of transmitted light (black and white) and autofluorescence in the Cy5 channel (scale in Fig 2A) at every time point. The time point is displayed on the top left and a 2 µm scale bar is displayed in the top right. A filled green arrow tracks a fluorescent punctum from time 4:20 to the end of the movie. An open green arrow tracks a fluorescent punctum that dissipates in a cell that ceases growth.

**Supplemental Movie 3. Δ*apcE* time-lapse with cell death and puncta dissipation replicate.**

Time-lapse fluorescence microscopy was performed to characterize the dynamics of the highly fluorescent puncta. Two colonies of Δ*apcE* was imaged every 20 minutes over 94 hours and 20 minutes. The movie displays a merged image of transmitted light (black and white) and autofluorescence in the Cy5 channel (scale in Fig 2A) at every time point. The time point is displayed on the top left and a 2 µm scale bar is displayed in the top right. A filled green arrow tracks a fluorescent punctum in the right colony for the entirety of the movie. An open green arrow tracks a fluorescent punctum that dissipates in a cell that ceases growth in the left colony.

**Supplemental Movie 4. Induction of *apcE* repression by CRISPRi.**

Time-lapse microscopy was performed to characterize the dynamics of highly fluorescent puncta induced in an *apcE* CRISPRi knockdown strain. Cells were precultured in the absence of aTC and IPTG so *apcE* expression was not repressed. IPTG and aTC were included in the agarose pad used during time-lapse microscopy to induce *apcE* knockdown. A single colony was imaged every 20 minutes over 24 hours and 40 minutes. The movie displays a merged image of transmitted light (black and white) and autofluorescence in the Cy5 channel (scale in Fig 2A) at every time point. The time point is displayed on the top left and a 2 µm scale bar is displayed in the top right.

**Supplemental Movie 5. Induction of *apcE* repression by CRISPRi replicate.**

Time-lapse microscopy was performed to characterize the dynamics of highly fluorescent puncta induced in an *apcE* CRISPRi knockdown strain. Cells were precultured in the absence of aTC and IPTG so *apcE* expression was not repressed. IPTG and aTC were included in the agarose pad used during time-lapse microscopy to induce *apcE* knockdown. A single colony was imaged every 20 minutes over 29 hours and 40 minutes. The movie displays a merged image of transmitted light (black and white) and autofluorescence in the Cy5 channel (scale in Fig 2A) at every time point. The time point is displayed on the top left and a 2 µm scale bar is displayed in the top right.

**Supplemental Movie 6. Recovery of *apcE* expression in *apcE* CRISPRi knockdown strain.**

Time-lapse microscopy was performed to characterize the dynamics of highly fluorescent puncta induced in an *apcE* CRISPRi knockdown strain. Cells were precultured with aTC and IPTG so *apcE* expression was impaired. Induction of *apcE* repression was removed through the exclusion of aTC and IPTG from the agarose pad used during time-lapse microscopy. A single colony was imaged every 20 minutes over 34 hours. The movie displays a merged image of transmitted light (black and white) and autofluorescence in the Cy5 channel (scale in Fig 2A) at every time point. The time point is displayed on the top left and a 2 µm scale bar is displayed in the top right. Different colored arrows (blue, red, and green) track puncta that dissipate over time until they are no longer visible.

**Supplemental Movie 7. Recovery of *apcE* expression in *apcE* CRISPRi knockdown strain replicate.** Time-lapse microscopy was performed to characterize the dynamics of highly fluorescent puncta induced in an *apcE* CRISPRi knockdown strain. Cells were precultured with aTC and IPTG so *apcE* expression was impaired. Induction of *apcE* repression was removed through the exclusion of aTC and IPTG from the agarose pad used during time-lapse microscopy. A single colony was imaged every 20 minutes over 35 hours and 40 minutes. The movie displays a merged image of transmitted light (black and white) and autofluorescence in the Cy5 channel (scale in Fig 2A) at every time point. The time point is displayed on the top left and a 2 µm scale bar is displayed in the top right. Different colored arrows (blue and red) track puncta that develop and dissipate during imaging.

